# From biting to engulfment: Target mechanics determines modes of phagocytosis through curvature–actin coupling

**DOI:** 10.64898/2026.01.28.702248

**Authors:** Shubhadeep Sadhukhan, Caitlin E. Cornell, Mansehaj Kaur Sandhu, Marta Batet Palau, Youri Peeters, Stijn Hanssen, Samo Penič, Aleš Iglič, Daan Vorselen, Daniel A. Fletcher, Valentin Jaumouillé, Verena Ruprecht, Nir S. Gov

## Abstract

Phagocytosis is a core innate immune process that clears targets spanning a wide range of mechanical properties, yet the role of target mechanics in recognition and engulfment remains unclear. Here, we combine theoretical modeling and experiments to reveal how target stiffness governs distinct modes of phagocyte–target interaction. We develop a membrane-based simulation framework in which both the engulfing cell and its target are deformable and undergo large shape changes, while actin-driven protrusions are regulated by curvature-sensitive membrane complexes. The model predicts three mechanical regimes with increasing target stiffness: (i) biting (trogocytosis), where part of the target is extracted; (ii) pushing, where the target is displaced rather than engulfed; and (iii) complete engulfment. We validate these predictions in epithelial clearance of apoptotic targets in vivo and macrophage engulfment of Giant Unilamellar Vesicles (GUVs) and lymphoma cells. Together, our results identify target mechanics as a key regulator of clearance and cell–cell interactions.

**Significance statement:** Phagocytosis is essential for immune defence, yet the physical principles governing engulfment of deformable targets remain poorly understood. Most theoretical models assume rigid particles, appropriate for phagocytosis of bacteria or fungi. When phagocytes engage dying cells or antibody-opsonised cancer cells, these targets undergo substantial shape changes during phagocytosis. We develop a theoretical model to simulate cell–cell interactions, enabling a mechanistic exploration of phagocytosis of soft targets. We utilize a model in which cytoskeletal protrusive activity is guided by curvature-sensitive membrane complexes, and show that target membrane rigidity dictates whether targets are fully engulfed, pushed away, or partially bitten. These mechanically driven dynamic regimes are validated experimentally using artificial elastic beads, GUVs, and lymphoma cells, both in vivo and in vitro.

## INTRODUCTION

Phagocytosis is essential for immune defense and tissue homeostasis, enabling cells to eliminate pathogens, apoptotic cells, and cancer cells through adhesion and engulfment [1, 2]. Although target recognition is primarily mediated by molecular signals, how target mechanics regulates phagocyte–target interactions remains poorly understood. Here, we combine theory and experiment to show that target stiffness is a key determinant of phagocytic behaviour.

Phagocytosis relies on dynamic actin recruitment to drive membrane protrusions that progressively wrap around the target [3–5]. However, how the actin recruitment and the resulting cytoskeletal force is coordinated with the membrane dynamics during this process is not well understood [6, 7]. We have recently shown that the coordinated recruitment of actin during phagocytosis of rigid objects can be explained very well by a theoretical model which includes a coupling between curved membrane protein complexes (CMC) that recruit and nucleate actin polymerization [8]. This model demonstrated that the presence of passive CMC can enhance the engulfment process, even in the absence of active forces. When active protrusive forces (which represent the protrusive forces due to actin polymerization) that are recruited by the CMC were considered, the model explained the more robust engulfment, at lower adhesion energies and less sensitive to the object’s shape [9] observed in cells [8]. By including the effects of active cytoskeletal forces, which are guided by the curvature coupling, this model goes beyond the theoretical descriptions of passive engulfment driven purely by adhesive forces [10–13]. Curved membrane proteins that recruit actin polymerization have been found experimentally to be associated with the leading edge of cellular protrusions [14, 15], which are involved in cellular adhesion, spreading and motility [16–18]. The theoretical model demonstrated that this curvature-actin activity coupling can explain many cellular shape and migration dynamics [19–21].

Motivated by these results, we explore here the interactions between a cell and a non-rigid object. We developed a minimal membrane-based model in which actin-driven protrusions emerge through curvature-sensitive membrane complexes and interact with deformable targets. We extend our model to allow for two vesicle-like surfaces to evolve and interact. For two symmetric vesicles containing passive CMC the model predicts a spontaneous symmetry-breaking transition, where one vesicle engulfs the other. For active CMC (describing the effects of actin polymerization in cells) interacting with a soft passive vesicle, the model predicts different dynamic regimes which depend on the rigidity of the target: with increasing rigidity, the active vesicle transitions from “biting”, to “pushing” and eventually “engulfing” the target. Despite its simplicity, the model predicts three distinct modes of phagocyte–target interaction that arise as a function of target stiffness: trogocytosis-like biting of soft targets, pushing of deformable targets without engulfment, and complete engulfment of stiff targets. These behaviors emerge from mechanical interactions alone, without invoking stiffness-dependent signalling pathways. These transitions arise in this model purely from the physical interactions and the curvature-force coupling.

To test these predictions, we examined phagocyte– target interactions across multiple experimental systems, including macrophage engulfment of Giant Unilamellar Vesicles and lymphoma cells, together with epithelial clearance of apoptotic targets *in vivo* using synthetic targets of defined stiffness. Across all systems, the observed behaviours closely matched the predicted stiffness-dependent transitions between biting, pushing, and engulfment. In particular, we validate the predicted intermediate “pushing” regime between trogocytosis and complete phagocytosis and show that it arises from curvature-dependent recruitment of actin at the cell membrane. These results identify a general physical mechanism by which target mechanics regulates phagocyte–target interactions across diverse cell and target types. Together, our findings establish target stiffness as a key regulator of cellular clearance during immune surveillance and tissue homeostasis.

### THEORETICAL MODEL

Our theoretical model is based on the Monte-Carlo (MC) calculation of the dynamics of a closed threedimensional triangulated self-avoiding vesicle with a spherical topology (Fig.1) [19, 22, 23] (See SI sections S1-S3 and Method and material for details). Within this model we denote the bare membrane nodes in blue, and the nodes containing the CMC in red. The active protrusive forces, representing the result of actin polymerization, are applied in the direction of the outwards local normal, at the locations of the CMC. In the present model we do not describe any contractile forces that can result from the activity of myosin-II motors. While myosin-II contractility affects the efficiency of phagocytosis and mostly gets activated during the late stages of the process, it is not an essential component [24–26]. These observations allow us to focus here on the regulation of the crucial recruitment of the actin cytoskeleton and its resultant protrusive forces, which dominate the phagocytosis process. Future extensions of our model will explore the role of contractile forces.

**FIG. 1.**
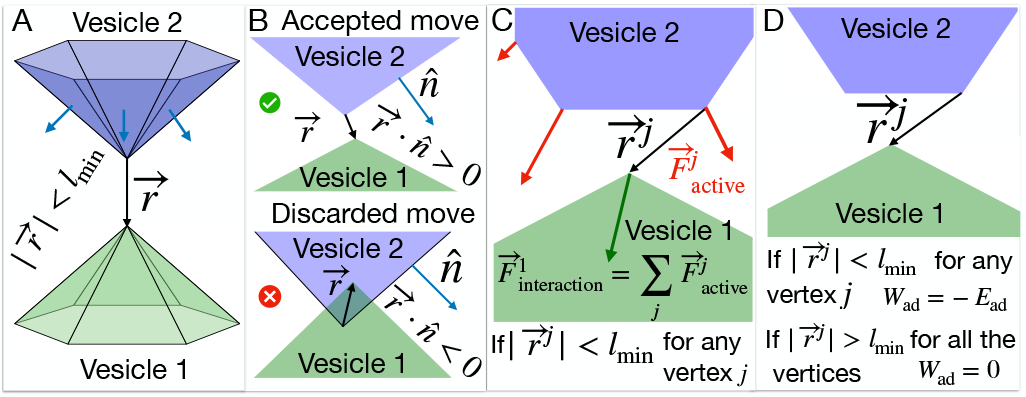
Interaction between two vesicles in the model: A) Two surfaces of two vesicles are shown in green and blue colors respectively. We check the distance between the vertices that belong to the two different vesicles and determine it they are interacting if the distance between them is less than the length unit of our simulation, i.e., *l*_min_. B) If the distance 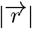 is less than *l*_min_, we find the dot product between the vector 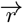 and the normals of all the triangles common to the vertex from the other vesicle 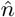 . If the dot product is negative, the MC move is discarded as the vesicles are overlapping; otherwise, it is accepted. C) The interaction force on the vertex of interest is the vector sum of the active forces applied by the vertices on the other vesicle within the interaction range. D) If a vertex from vesicle 1 is at a distance less than the interaction range *l*_min_ from any vertex of another vesicle, then the adhesive energy between them is −*E*_ad_ for both of these vertices. Otherwise, it is zero.

Our model is purely a membrane model, and it does not include any information about the details of the actin network inside the cell or the internal structure of the cell, such as organelles (nucleus) or bulk elastic deformations and their associated energy cost.

Here we developed our previous model to allow for the interaction between two dynamic vesicles (Fig.1). As a first step, we find the nodes that are adjacent between the two vesicles (Fig.1A). Such proximal nodes are restricted in their MC moves, as we do not allow the two vesicles to pass through each other. We need to check any such pair of vertices from different vesicles (Fig.1B, Fig.S1).

Next, we consider the active forces that a vesicle exerts on its neighboring vesicle. Any vertex in vesicle 1 feels the active force due to the other vesicle’s active CMC sites, that are within the interaction range (Fig.1C). The effect of this active force is included by adding the energy term given by,

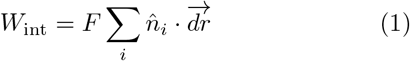

where *i* runs through all the vertices belonging to vesicle 2 within the distance of *l*_min_ from the vertex of interest in vesicle 1, the force vectors have amplitude *F* and are directed at the outwards normal 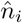 at the sites *i*, while 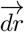 is the MC move of the vertex in vesicle 1. Note that in our model we do not explicitly maintain force balance. When adhered to external surfaces, they serve as momentum sinks, while for free vesicles we work in the center-of-mass frame which allows us to calculate relative shape changes, such that globally the force is effectively balanced.

Finally, we introduce an adhesion energy between proximal vertices on the two neighboring vesicles. Each vertex *j* that is within the adhesion range to the other vesicle (Fig.1D) has an adhesion energy that is given by,

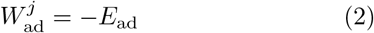

Initially, we evolve it for a few Monte-Carlo steps from a pentagonal-dipyramid, until it becomes nearly spherical. The active force (when implemented) is set to *F* = 2 *k*_*B*_*T/l*_min_ and the protein-protein interaction energy is set to *w* = 1 *k*_*B*_*T* throughout the paper.

The area of the vesicles in our model is roughly conserved, with the bond lengths limited in their allowed range of fluctuation (*l*_min_ *< l <* 1.7 *l*_min_) [22]. The typical changes in area in our simulations (without applied osmotic pressure) are of order 5 % (see Fig.S7). When the osmotic pressure is applied inside the vesicle, the bond lengths are pushed towards their maximal allowed value (and therefore the average area increases), and the fluctuations in area are greatly suppressed. Finally, we note that the MC method does not include dissipative processes such as membrane-membrane and viscous friction, and therefore do not give information about the physical time-scale of the simulated process.

## RESULTS

### Spontaneous symmetry breaking of adhering vesicles containing CMC

We first analyzed the adhesion between two identical vesicles, in the absence of active forces (*F* = 0). For calibration purposes, we tested our model for adhesion of bare membrane vesicles (no CMC) and under the conditions of volume conservation (Figs.S2-S4, See SI section S6), for which analytic solutions are available [27]. In this limit, we could extract the contact angles between the two vesicles, and compare them to an analytic result [28–30]. The agreement between the simulations and the analytic calculation serves to validate our numerical procedure. Adding CMC induces larger spreading and some breaking of the rotational symmetry, though the two adhering vesicles remain symmetric with respect to each other (Fig.S4). In the presence of CMC the contact surface between the vesicles is not anymore composed of flat or spherical surfaces, as was found for simple adhered vesicles [27, 31].

When we remove the volume conservation constraint (See SI section S7), we find that the presence of CMC can drive a spontaneous symmetry breaking above a critical value of the adhesion energy (Fig.2A). At low adhesion strength the two vesicles are still symmetric (Fig.2A, *E*_ad_ = 1 *k*_*B*_*T* ), but above a critical adhesion strength one of the vesicles spontaneously forms a cup-like shape (top vesicle in Fig.2A, vesicle 2), which partially encapsulates the other vesicle (bottom vesicle in Fig.2A, vesicle 1). The CMC in the top vesicle condense along the sharp rim of the cup shape, similar to the organization of such a vesicle when engulfing a rigid sphere [8] (Fig.3A). The bottom vesicle remains largely spherical, with the CMC randomly spread as small isolated clusters. This transition is driven by a lowering of the total energy of the two vesicles. The bending energy increases during the symmetry-breaking transition (Fig.S5), as the cup-shaped vesicle 2 is highly curved. However, this increase is offset by the adhesion energy between the vesicles which decreases the total energy (Fig.2B-E), and to a much smaller amount by the CMC-CMC binding energy (Fig.S5). The huge changes to the volume of the cup-shaped vesicle 2 during this spontaneous shape transition are quantified in Fig.2F,G.

**FIG. 2.**
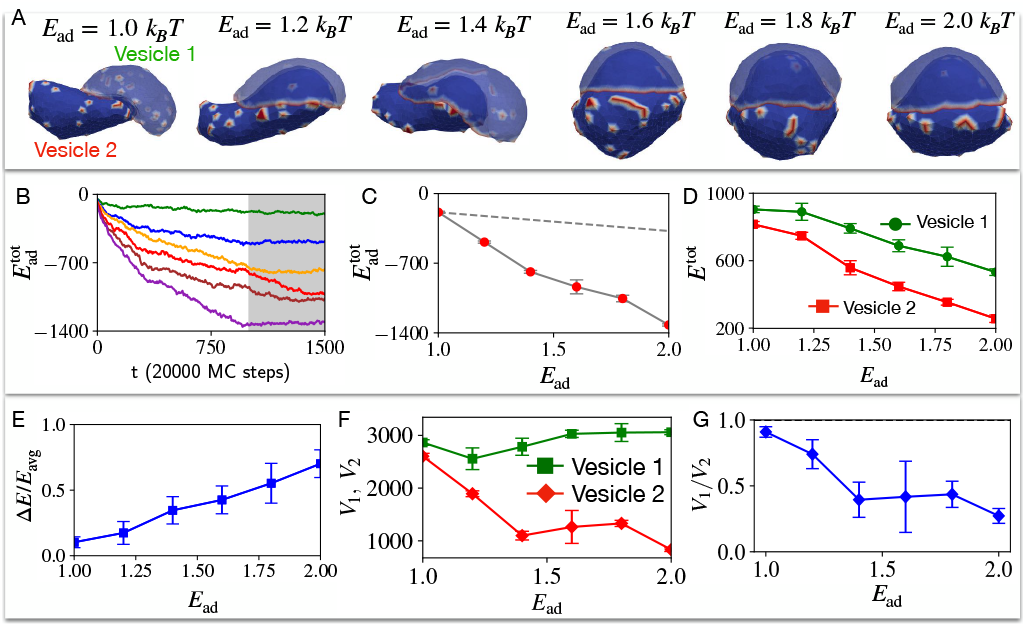
Two identical vesicles adhere to each other for different strengths of adhesion energy parameter *E*_ad_. A) Final configuration snapshots of the pair of passive vesicles (at time step *t* = 1500), for the adhesion energy parameter *E*_ad_ = 1, 1.2, 1.4, 1.6, 1.8, 2.0 in units of *k*_*B*_ *T* . Blue denotes the bare membrane nodes, while red at the passive CMC nodes. B) The average adhesive energy 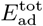 per vesicle is shown as a function of timestep for *E*_ad_ = 1, 1.2, 1.4, 1.6, 1.8, 2.0 in units of *k*_*B*_ *T* with green, blue, orange, red, brown and purple solid lines respectively. C) The final total adhesive energy 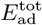 (averaged over the grey shaded time window shown in (B)) is shown for six different values of *E*_ad_. The grey dashed line represents the total adhesive area in the case of *E*_ad_ = 1 *k*_*B*_*T* multiplied by the *E*ad. It shows that the total adhesive energy increases due to the increase in adhered area, faster than the increase in *E*_ad_ (dashed line). In D) we show the total energies the pair of vesicles, averaged over the grey shaded region for six different values of *E*_ad_. E) The relative difference between the vesicles increases with *E*_ad_. F) Average volumes of the vesicles as function of *E*_ad_, and G) the corresponding volume ratio. We used 722 vertices for each vesicle of bending rigidity *κ* = 20 *k*_*B*_ *T* , out of which 50 vertices represent the curved-protein complexes with intrinsic curvature 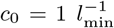 i.e. the CMC percentage is *ρ* = 6.93 %. Here, volume is not conserved.

**FIG. 3.**
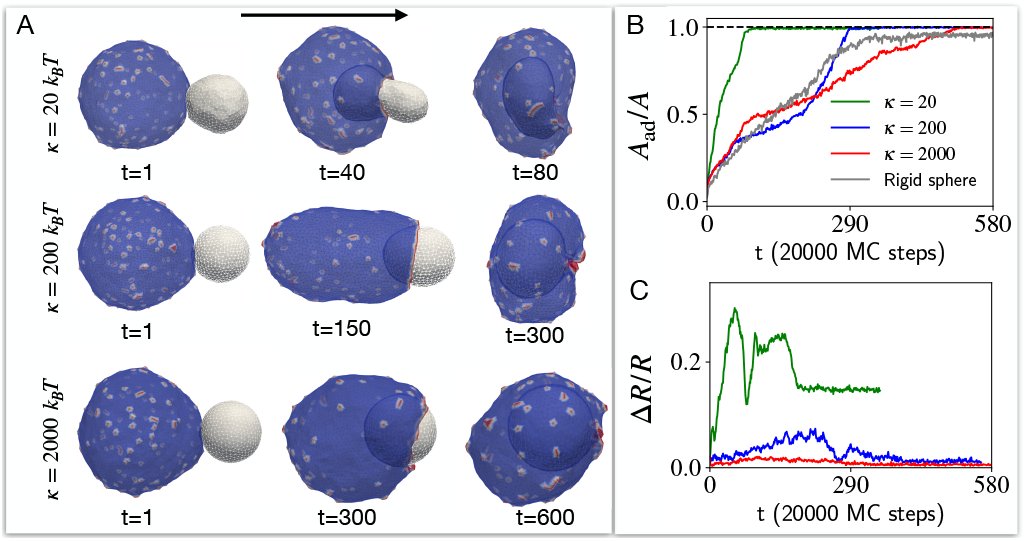
The engulfment of target vesicles of different rigidities by a cell-like vesicle with passive CMC. A) The snapshots of the shapes of the interacting vesicles with time for different bending rigidity *κ* values of the target vesicle. We set the bending rigidity of the cell-like vesicle at 20 *k*_*B*_ *T* . B) The time evolution of adhered area fraction of the target vesicle for different *κ* values (see SI section S4 for details on this calculation). As *κ* increases, the outcome is approaching the completely rigid *κ* =*∞* limit that is shown in grey. C) Time evolution of the deviation of the target vesicle from a spherical shape, for different values of *κ* (color code as in (B)), measured by the relative standard deviation for the position vector of all the vertices with respect to the center of mass of the vesicle (Eq.S11). As *κ* increases, the target remains spherical all the time, and it takes longer time to engulf as it requires the cell-like vesicle to extend its adhesion cup over a bigger crosssection.

Shapes that are similar to the vesicle pair of Fig.2(A) were obtained for adhering soft tissues (modeled as vesicles), where the symmetry breaking was induced by a large difference in the active surface tension between the two vesicles [32]. In our system the symmetry breaking is spontaneous, and the two vesicles have identical properties.

### Rigidity-dependent biting, pushing and engulfing

Next we explore the process of engulfment that mimics the phagocytosis of a non-rigid object by a cell. The engulfed (target) object is described by a small vesicle (*N*^*T*^ = 847 vertices, forming a spherical object with radius of 10 *l*_min_), and the cell-like (bigger) vesicle has 3127 vertices. We set the CMC concentration *ρ* = 4.8 % on the cell-like vesicle throughout this work. We start by validating our computation, demonstrating that for a very rigid target vesicle (bending modulus *κ* = 2000 *k*_*B*_*T* ) the engulfment proceeds in the same manner as we previously computed for a perfectly rigid sphere (Fig.S6, and see SI section S8) [8]. Note that we are using the bending modulus as a model parameter to easily control the rigidity of the target vesicle, with a higher bending modulus diminishing shape deformations that increase the local mean curvature.

We start exploring how a cell-like vesicle that contains passive CMCs (*F* = 0, Fig.3A), engulfs a target vesicle of different bending modulus. We find that the cell-like vesicle with passive CMC is able to fully engulf the target vesicle, with the engulfment proceeding faster for the softest target vesicle (See Movie S1). This faster engulfment is facilitated by the large deformation (See SI section S9 for details of deformation measurement) of the target vesicle (as shown in Fig.3B,C), which enables the cell-like vesicle to extend an adhesion cup over a smaller cross-sectional area.

This behaviour is drastically changed when the CMCs induce active protrusive forces. We explore in Fig.4 the engulfment dynamics as function of the bending modulus *κ* of the target vesicle, and find three main dynamical phases. We start with the high *κ* regime (blue traces in Fig.4A, and blue region on the phase diagram Fig.4B), where the cell-like vesicle completely engulfs the rigid target vesicle (similar to the engulfment of a rigid object, Fig.S6 [8]), as shown in the snapshots of Fig.4E. For a softer target vesicle (green traces in Fig.4A, and green region on the phase diagram Fig.4B), we find that the target vesicle ends up being pushed away (Fig.4D), and the contact area stalls in a “suction-cup”-like shape, and later retracts (Fig.4A) until the two vesicles detach (zero final adhered area fraction, Fig.4B) due to the high bending energy of the elongated “finger” attached to the target vesicle (Fig.S8, See SI section S10). At even lower values of *κ* (red traces in Fig.4A, and red region on the phase diagram Fig.4B) we find that the engulfed area stalls at a small value (*A*_*ad*_*/A <* 0.5, Fig.4A,B), corresponding to a “biting”-type dynamics (Fig.4C). Note that we do not allow the vesicles to undergo fission, even when greatly deformed.

**FIG. 4.**
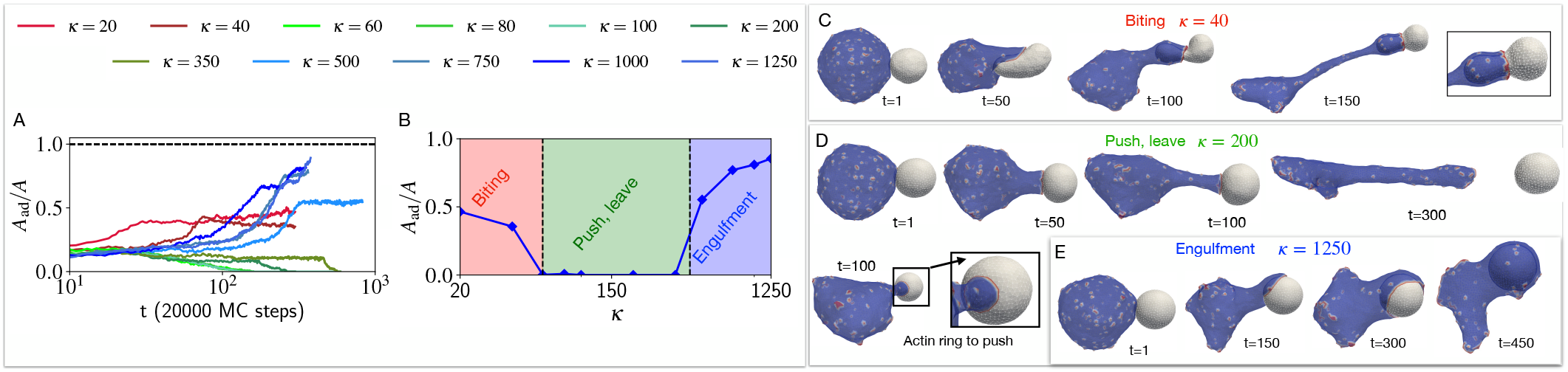
Effect of the bending rigidity of the target vesicle on the process of phagocytosis. The time evolution of the adhered area fraction of the target cell is shown in (A), for different bending modulus values. B) The final adhered area fraction of the target vesicle is shown as a function of the bending rigidity *κ*. We indicated three phases of biting, push-leave, and engulfment with red green and blue as *κ* increases. The time evolution of the shapes and the snapshots are shown for three example of biting, push-leave, and engulfment in C), D) and E) panels, where the bending rigidity *κ* for the target vesicle is set to 40 *k*_*B*_*T*, 200 *k*_*B*_*T*, and 1250 *k*_*B*_*T* respectively. We set the bending rigidity of the cell-like vesicle to 20 *k*_*BT*_.

The origin of these dynamical phases can be understood when we investigate how the membrane shape and the orientation of the active forces are coupled. In Fig.5(A) we define the components of the active force that is exerted by a CMC of the cell-like vesicle when its in contact with the nodes of the target vesicle. This force is applied towards the outwards normal of the CMC, and has both normal (*F*_⊥_) and tangential (*F*_∥_) components with respect to the target vesicle surface.

**FIG. 5.**
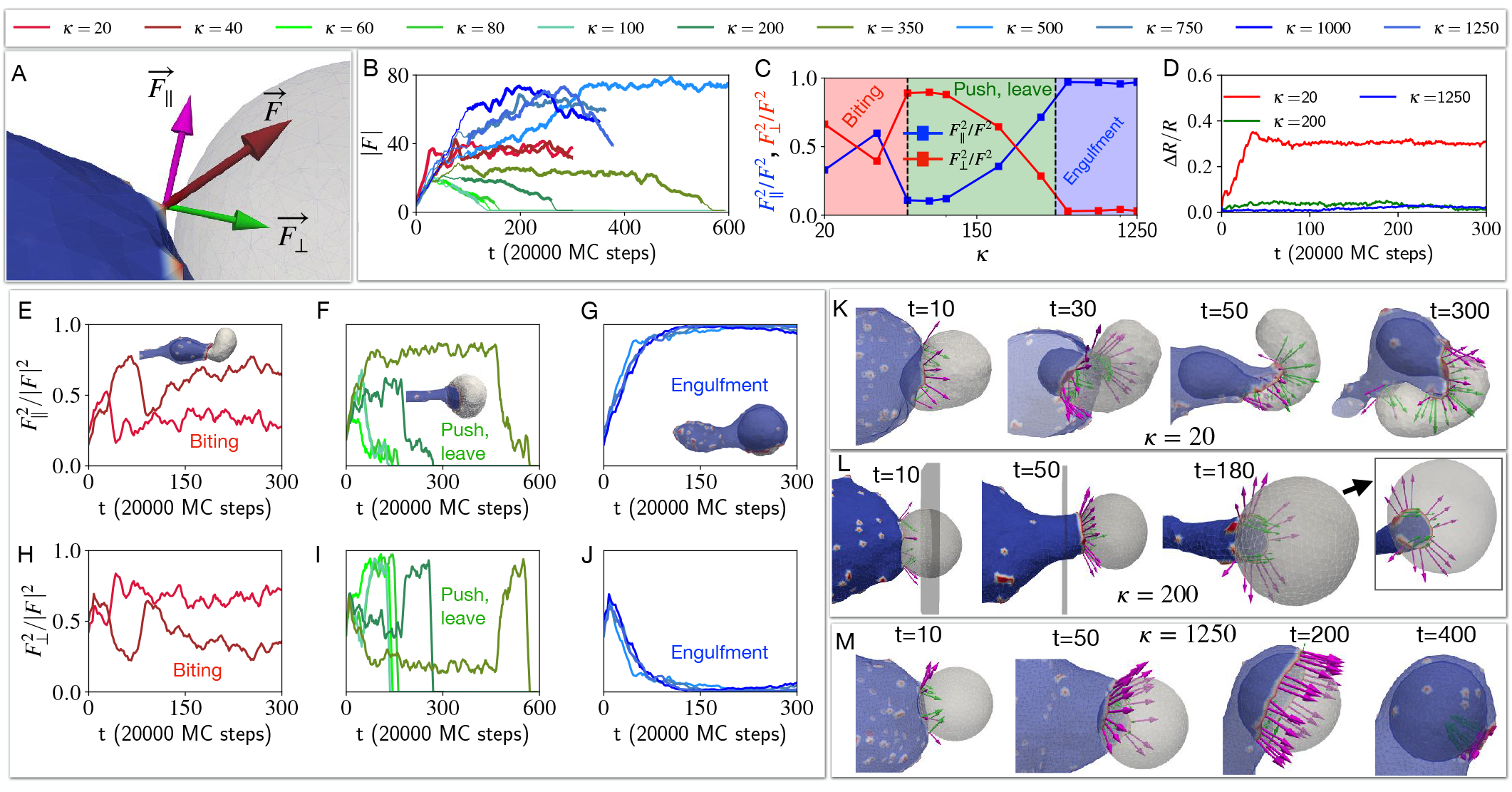
Coupling between the target vesicle deformation and alignment of the active forces during the engulfment process. A) The active force due to an active CMC node of the cell-like vesicle is applied to a neighboring node of the target vesicle 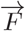 . This force is decomposed into two parts, 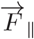 is tangential to the surface of the target vesicle and 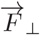 is the pushing normal force (shown in magenta and green colour, respectively). B) The time evolution of the total magnitude of the force applied to the target vesicle by the cell-like vesicle is shown, for different bending rigidities of the target vesicle (as in Fig.4). C) Time averaged the tangential and normal force fractions as function of the bending rigidity of the target vesicle *κ* (over the times denoted by bold lines in (B)). D) The deviation of the target vesicle from a sphere is shown for three different bending rigidities *κ* of the target vesicle (Eq.S11). E)-G) show the fraction of tangential force on the target in the three dynamical regimes of biting, push-leave, and engulfment. H)-j) Similarly, the fraction of normal pushing force on the target in the three dynamical phases. K)-M) Snapshots showing the force decomposed into tangential and normal components together with the deformations of the target vesicle. We show how the early deformation caused by the normal components affect the later alignment of the CMC at the leading edge of the adhesion patch. K) A very soft target vesicle can initially adhere and bend into the cell-like vesicle, during the early stages. However, the CMC then impinge against the remaining target vesicle, and end up pushing and twisting it with a significant normal component. L) In the pushing regime the deformation of the target vesicle is sufficient to prevent the CMC from aligning tangentially, and a significant normal component maintains the pushing dynamics. M) For the rigid target vesicle the CMC cluster aligns tangentially and drives efficient engulfment. The bending rigidity of the bigger cell is set to 20 *k*_*B T*_

During the engulfment phase of rigid target vesicles we find that the CMC form large leading-edge clusters (large value of the total transmitted force, blue traces in Fig.5B) which mostly exert tangential forces (Fig.5G,J). This is shown in the snapshots of Fig.5M. The average fraction of the two force components (beyond the transient initial cup formation stage, denoted by the bold lines in Fig.5B), is shown in Fig.5C, and clearly demonstrate the relation between these force components and the resulting three dynamical phases.

In the pushing phase of slightly softer target vesicles we find that the leading-edge cluster gets arrested at a smaller size (green traces in Fig.5B), eventually disappearing when the two vesicles disengage. The dynamics of the force components (Fig.5F,I) show that the normal component remains relatively high, preventing the efficient spreading of the engulfing membrane over the target vesicle. In Fig.5L the snapshots show the origin of this behavior: due to the deformation of the target vesicle, the leading-edge of the engulfing membrane is not tangentially oriented, further pushing into the target vesicle and maintaining its deformation. This feedback between shape and force orientation arrests the spreading, as the target vesicle is pushed and deformed.

In the regime of softest target vesicle (red traces in Fig.5B) the target vesicle gets strongly deformed by the adhesion to and active forces exerted by the engulfing membrane (red line in Fig.5D). This large deformation prevents the CMC and the active force from aligning tangentially, with both components having similar magnitude (Fig.5E,H). Due to the large deformations of the soft target vesicle (Fig.5K) the leading-edge ends up mostly pushing it after a small portion is engulfed, leading to a “biting”-like behavior. This is very different from the smooth engulfment observed when the active forces are absent (Fig.3).

In Fig.S9 we show the same set of dynamic phases when we vary the osmotic pressure (see SI section S12 for details) inside the target vesicle (while keeping *κ* = 20 *k*_*B*_*T* ). We find that as the internal pressure *p* inside the target vesicle increases, the membrane of this vesicle gets stretched out and tense, inhibiting any shape changes and effectively stiffening the vesicle. Not surprisingly, the observed phases, as function of the pressure *p* (Fig.S9), perfectly match the phases observed as function of bending rigidity in Fig.4.

To summarize, in our model the recruitment and organization of the actin-derived forces at the leading edge of the cell are curvature-sensitive. This curvature-activity coupling makes the forces’ alignment dependent on the target deformation, and gives rise to three dynamical phases as function of the target’s rigidity. These theoretical predictions, especially the emergence of an intermediate “pushing” phase, will next be compared to experimental observations.

### Comparison to experiments

We start by validating our model for the localization of actin polymerization at the leading edge of the phagocytic cup, and how it transmits forces to the engulfed object. We compare to experiments utilizing elastic beads [33, 34], which enable the extraction of the forces exerted by the cell during the phagocytosis process (Fig.S12). Since the beads are composed of a cross-linked gel that has bulk elasticity, we roughly emulate it by adding an effective spring network to our membrane (details in SI section S13). This is a rough approximation, compared to detailed theoretical descriptions of cross-linked gels engulfed by membranes [35, 36]. Nevertheless, the comparison shows very good overall similarity in both the deformations of the beads, and the associated localization of the normal forces exerted by the cell’s leading edge. This good agreement validates that protrusive forces in the model are exerted realistically both spatially and temporally, supporting our association of the actin force with the highly curved CMC cluster in the simulations. A band of normal force exerted by this leading edge on the engulfed sphere results in local squeezing of the sphere, in both experiments and simulations. We also observe that the actin at the leading edge is non-uniform and fragmented in both the experiments and our simulations, suggesting that a simple coupling between curvature and actin nucleation may be involved in a complex and nonuniform organization of this moving front [37, 38] due to its sensitive dependence on the target deformations.

Following the validation of the forces exerted in our model, we now compare to experimental data showing the dependence of the phagocytosis dynamics on target stiffness. Our model’s prediction of a “pushing” phase for intermediate target stiffness offers an explanation to the puzzling observations of cells either engulfing or pushing away apoptotic cells during embryogenesis [39]. To directly test this prediction of our model that the mechanical state of apoptotic targets determines whether engulfment or pushing behaviors take place, we used the early zebrafish embryo as an in vivo model system. Epithelial cells in the early embryo have previously been shown to generate two distinct types of protrusions: phagocytic cups that mediate apoptotic cell engulfment and “epithelial arms” that exert pushing forces and promote apoptotic cell displacement [39]. We therefore aimed to assess if target stiffness may represent a key biophysical cue governing the choice between these two modes of phagocyte-target interaction. For this purpose, we employed synthetic apoptotic targets with tuneable mechanical properties that recapitulate the lipid composition and phosphatidylserine (PS) exposure of apoptotic cells (termed lipobeads in the following). Using quantitative live in vivo imaging, we first confirmed that soft lipobads (1.3 kPa) versus stiff lipobeads (80 kPa) were efficiently recognized and internalized by epithelial cells (Fig. S11A,D). High-resolution imaging further captured individual uptake events, demonstrating bona fide phagocytic engulfment of lipobeads by the epithelial tissue (Fig. S11B,C).

Notably, whereas stiff lipobeads (80 kPa) were exclusively engulfed by epithelial cells (Fig. S11A), soft lipobeads (1.3 kPa) exhibited two distinct outcomes: they were either actively displaced by epithelial cells through pushing interactions (Fig. 6A) or subsequently engulfed (Fig. 6B). Consistent with these observations, quantitative tracking revealed a pronounced increase in target mobility for soft compared with stiff lipobeads (Fig. 6C,D; Movie S12). Apoptotic cells generated through overexpression of the pro-apoptotic factor Bax displayed the highest degree of mobility, in agreement with a softening of apoptotic cells [40, 41]. Together, these results provide direct in vivo evidence that apoptotic target stiffness is a major determinant of epithelial protrusion dynamics and thus clearance behaviour. Soft lipobeads and apoptotic cells were frequently observed to be displaced by epithelial cells and subsequently detach from the pushing cell (Fig. 6A)[39], as predicted by our simulations (Fig. 4D).

**FIG. 6.**
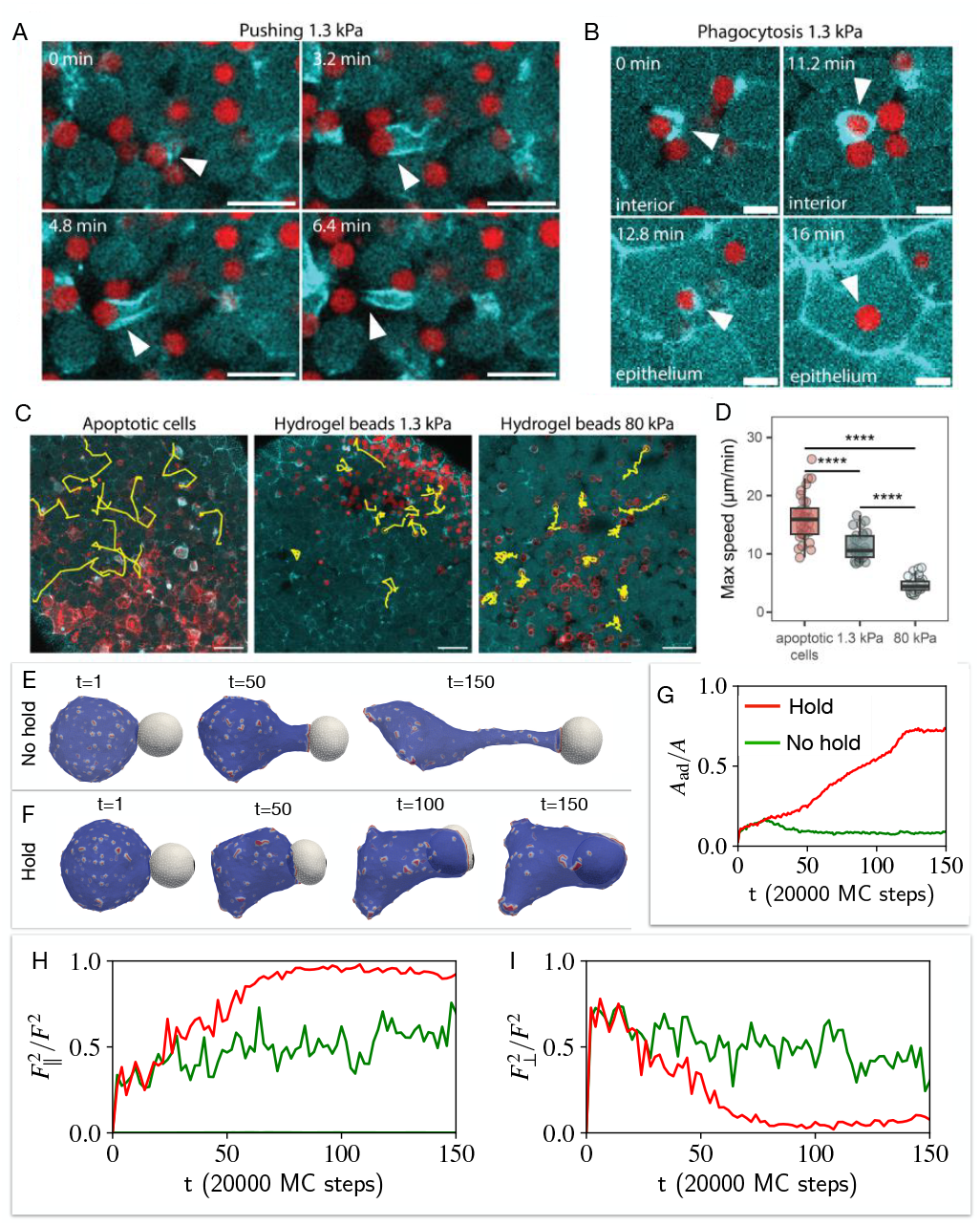
Epithelial phagocytosis and mobility of apoptotic cells and synthetic targets with variable target stiffness in vivo. A. Representative images of a pushing event of 1.3 kPa hydrogel beads over the indicated time period in vivo. Arrowheads point at the formation of an epithelial pushing protrusion termed ‘epithelial arm’, which leads to the movement of the synthetic apoptotic target in vivo. B. Representative images of an individual phagocytosis event (arrowhead) of 1.3 kPa hydrogel beads in vivo. C. Representative x/y-trajectories (yellow lines) obtained from 3D tracking of apoptotic cells (Bax+ cells, red, left), 1.3 kPa hydrogel beads (red, middle); and 80 kPa hydrogel beads (red, right) in embryos expressing Lifeact-GFP (cyan). D. Analysis of maximum target speed for apoptotic cells (red, n = 32 tracks from 3 embryos), 1.3 kPa hydrogel beads (dark cyan, n = 29 tracks from 3 embryos) and 80 kPa hydrogel beads (light cyan, n = 30 tracks from 3 embryos). Data points represent individual tracks. Pairwise comparisons using Wilcoxon rank sum test *p <* 0.0001 between all conditions. All embryos were obtained from the Tg(actb1:Lifeact-GFP) line. Size of hydrogel beads: 8.8*μ*m (1.3 kPa), 8.2*μ*m (80 kPa). E. Shows the snapshots of cell-like vesicle pushes the target away when the target is not held by any external means. F. Shows the snapshots of how the cell-like vesicle starts to engulf the target while it is held by some external means (See SI section S11). H. The fraction of force applied by the cell-like vesicle parallel to the target’s surface. I. The fraction of force applied by the cell-like vesicle normal to the target’s surface. Scale bars: 20*μ*m (A), 10*μ*m(B), 40*μ*m (C).

Engulfment of soft lipobeads was observed when the target became mechanically constrained by neighbouring epithelial cells (Fig. 6B), suggesting that local tissue geometry can modulate the outcome of cell–target interactions. To investigate the role of spatial constraints, we incorporated target confinement into our model by fixing a small region of the target vesicle in space (black patch at the pole opposite to where the cell is at t=1, Fig. 6E,F). This constraint has a substantial effect on phagocyte-target interaction, resulting in engulfment (Fig.6G), as it allows the protrusive forces at the leading edge of the cell-like vesicle to align tangentially to the target (Fig.6H,I), which is needed to drive efficient engulfment. In contrast, unconstrained soft targets are displaced before stable wrapping can be established, resulting in pushing-mediated target transport rather than uptake.

In Fig.7 we compare the theoretical predictions of three dynamical phases as function of the membrane tension of the target vesicle, to experimental observations. Fig.7(A-C) gives examples of snapshots of the interactions between macrophage cells (labeled in green) and giant unilamellar vesicles that contain antibodies that trigger macrophage adhesion (GUVs, labeled in pink) [42]. In the experiments the membrane tension was varied using different sucrose solutions to fill the GUVs, and as function of decreasing membrane tension the observed behavior changes from (mostly) engulfment at high tension (Fig.7A), to a mixture of pushing and some biting at intermediate tensions (Fig.7B), finally exhibiting biting (trogocytosis) activity for the lowest GUV membrane tensions (Fig.7C, See Movies S2, S3, S4).

**FIG. 7.**
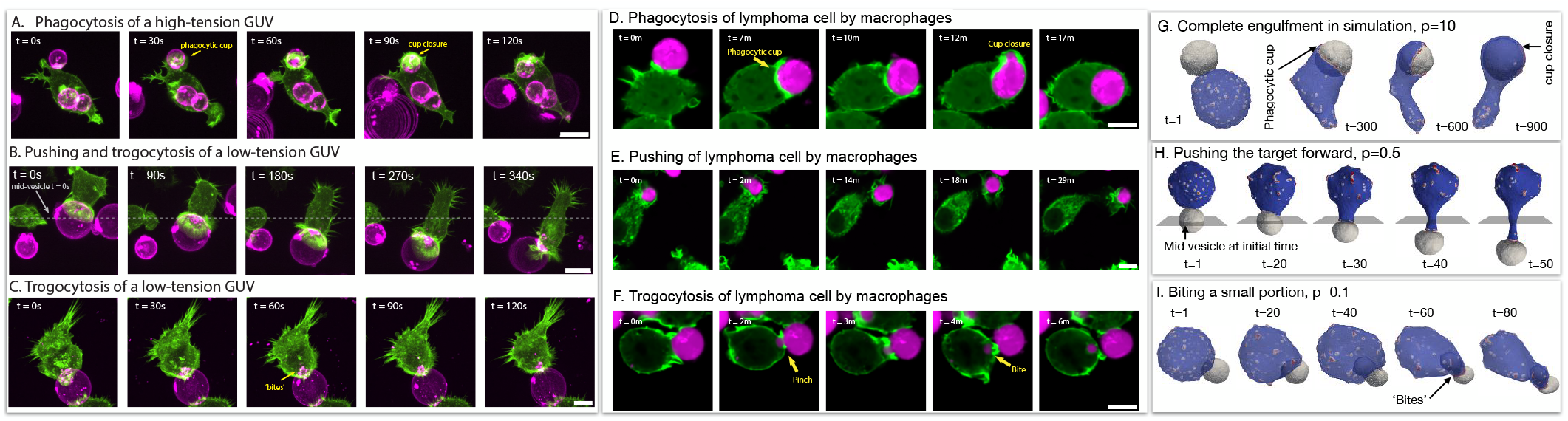
(A) Giant unilamellar vesicles (GUVs) composed of POPC and biotin-DOPE and filled with a hyperosmotic sucrose solution (300 mOsm), leading to taut high-tension vesicles, are opsonized with AlexaFluor647 anti-biotin and mixed with LifeAct-GFP-expressing macrophages. During phagocytosis, the macrophage forms a phagocytic cup, identifiable from enriched actin in a ring, that encircles the GUV. The macrophage fully engulfs the GUV within tens of seconds. (B) GUVs filled with a hypoosmotic sucrose solution (270 mOsm) are at low tension and can be ‘pushed’ by a macrophage, as shown by the position of the vesicle over time relative to the mid-plane at t=0s. (C). When low-tension GUVs are trogocytosed by a macrophage, punctate ‘bites’ can be observed within the macrophage. Scale bar is 5 μm. (D-F) Time-lapse confocal microscopy images show interactions between macrophages (Lifeact-mScarlet3, pseudo-colored in green) and antibody-opsonized lymphoma cells (pseudo-colored in magenta). D) Example of a macrophage engulfing an entire target cell. E) Example of a macrophage pushing the target cell. F) Example of a macrophages trogocytosing a fragment of the target cell. Scale bars are 10 *μ*m. In simulations, (G) the complete engulfment of the target vesicle when its osmotic pressure is high 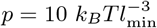 . (H) The target is pushed when the osmotic pressure is intermediate 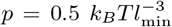 A plane that is shown perpendicular to the pushing direction through the middle of the target vesicle’s initial position. (I) For a very low osmotic pressure, 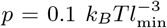 the cell-like vesicle takes a bite from the target.

Our model and observations suggest that the engagement of target populations that display heterogeneous stiffnesses, such as cancer cells, could lead to a mix of pushing, partial and complete phagocytosis. To assess this possibility with physiological targets, we performed time-lapse microscopy of macrophages (green) interacting with lymphoma cells opsonized with therapeutic antibodies (magenta) (Fig.7D-F). From top to bottom we see the same three typical behaviors: engulfment (Fig.7D), pushing (Fig.7E) and biting (trogocytosis, Fig.7F) (See Movies S5, S6, S7 respectively). While we do not know exactly the stiffness of each of these cancer cells, finding this range of behaviors and comparing to our model suggests that cancer cells exhibit a wide range of stiffness [43]. This trait of cancer cells was indeed measured using Atomic Force Microscopy (AFM) [44, 45].

In Fig.7(G-I) we demonstrate how the three phases observed in the experiments are predicted by our theoretical simulations as a function of the pressure inside the engulfed target vesicle (See Movies S8, S9, S10, respectively). Higher internal pressure acts to suppress dynamic changes and fluctuations in the target vesicle’s area, equivalent to higher membrane tension (Fig.S10). Note that we do not allow our vesicles to undergo topological changes such as fission, so the biting behaviour in the simulations is arrested.

## DISCUSSION

Our combined experimental and theoretical study identifies target mechanics as a previously underappreciated regulator of phagocyte behaviour, with relevance for the clearance of opsonized targets, pathogens and apoptotic cells. While target recognition and uptake has primarily been addressed from molecular receptorligand signalling, our findings strengthen increasing evidence that the mechanical target state provides critical information that influences the mode of a phagocyte’s response.

We have explored here the dynamics of cellular adhesion and engulfment of soft objects, whereby the mechanism of the self-organization of the protrusive forces of actin polymerization is through curved membrane protein complexes (CMC). Curved membrane proteins that recruit actin polymerization were shown to drive cellular protrusions during cell adhesion and migration [16, 18, 19]. This mechanism was previously shown to explain how cells phagocytose rigid objects [8], and here we explored this mechanism when the engulfed object is flexible.

Our theoretical model of phagocytosis goes beyond passive models of engulfment driven purely by adhesion [10], by describing the role of active cytoskeletal forces. A previous work that included active cytoskeletal forces [46]also found engulfment rate that diminishes when the target is softer, due to the target deformation affecting the cell’s ability to spread by adhesion and random active force. In our model we provide an explicit mechanism that explains how the target deformation affects the organization of the cell’s cytoskeleton, by coupling the recruitment of actin polymerization to the shape of the cell and the target, through curvature-sensitive membrane protein complexes.

Our model predicts that as function of the stiffness or membrane tension of the engulfed vesicle, there are distinct dynamical phases: While a stiff object is engulfed, a softer object will be spontaneously pushed away until it detaches. The softest objects get partially engulfed, which we expect in reality to result in a piece getting “bitten” off (trogocytosis). The theoretical model explains the origin of these dynamical phases as arising due to the feedback between the deformations of the engulfed object, and the orientation of the active forces exerted by the cell’s leading edge, due to the curvature-activity coupling. Comparing to experiments of in-vivo epithelial phagocytosis of apoptotic cells and artificial beads, as well as in-vitro immune cells engulfing artificial vesicles or cancer cells, we validate the predicted relation between engulfment dynamics and the mechanical deformability of the target. Crucially, the experiments validate the theoretically calculated “pushing” phase, which was predicted and observed to emerge at intermediate target stiffness.

Here we show that epithelial cells selectively push soft versus stiff targets prior to engulfment. In the context of apoptotic target clearance by epithelial cells, this increased pushing of apoptotic targets thereby facilitates a cooperative clearance that reduces the overall time it takes for epithelial cells to remove apoptotic cells from the tissue [39]. Furthermore, our data suggest that local tissue geometry can modulate these mechanically driven outcomes, as confinement by neighbouring cells promotes engulfment of otherwise displaceable soft targets. The observed dependence of phagocyte dynamics on target stiffness is particularly intriguing considering mechanical changes that accompany apoptosis. During apoptosis progression, mechanical structures within the cell are modified, leading to changes in cortical tension, cytoskeletal organization and cellular stiffness [40, 41]. Our results raise the possibility that such mechanical remodelling is not merely a consequence of cell death but can actively influence how dying cells are handled by phagocytic cells.

Our theoretical results offer an explanation of the lower rates of successful engulfment observed for soft beads [34, 47, 48], in agreement with our simulation results (Fig.4). Similarly, red-blood cells were observed to be phagocytosed more easily, and even overriding the “self” - signal, when rigidified [24]. The model results give insight into the observed dependence of engulfment and trogocytosis on cell membrane tension [49] and rigidity [50].

Note that our model predicts that successful engulfment of soft targets is facilitated by the cell employing weaker cytoskeletal forces (Fig.3). Altogether, these results suggest possible future interventions to enhance or inhibit phagocytosis and trogocytosis based on the interplay between the target’s stiffness and the protrusive activity of the engulfing cell.

Our model is using only the most minimal physical components and forces, and therefore offers a path to obtaining deep and general understanding of an important biological process which is shared by many cell types [51– 53]. Future extensions of our modeling approach could include additional processes, such as myosin-induced contractility and the effects of actin treadmilling-induced forces, in addition to more complex description of the adhesion between the cell and its target (which may itself depend on stiffness and applied forces [42]).

### I. METHODS AND MATERIAL

#### GUV electroformation

Solutions containing 0.25 mg total lipids were spread evenly on slides coated with indium tin oxide (70-100 Ω/sq; Sigma Aldrich). The slides were placed under vacuum for >30 min to allow for complete evaporation of chloroform. A capacitor was created by sandwiching a 0.3-mm rubber septum between two lipid-coated slides. The gap was filled with 200 *μ*L of 300 mM sucrose (hyperosmotic solution compared to PBS (285 mOsm) to make high-tension GUVs) or 270 mM sucrose (hypoosmotic solution compared to PBS to make low-tension GUVs). Sucrose solution osmolarity was measured using an osmometer (Precision Systems). GUVs 10 to 100 *μ*m in diameter were electroformed by application of an AC voltage of 1.5 V at 10 Hz across the capacitor for 1 h at 55°C.

#### Imaging techniques

All live cells were maintained at 37°C, 5 % *CO*_2_ with a stage top incubator (Okolab) during imaging. For confocal microscopy, cells were imaged with a spinning disk confocal microscope (Eclipse Ti, Nikon) with a spinning disk (Yokogawa CSU-X, Andor), sCMOS camera (Prime 95B, Photometrics), and a 60x objective (Apo TIRF, 1.49NA, oil, Nikon). The spinning disk confocal microscope was controlled with Nikon Elements (Nikon). Images were analyzed and prepared using FIJI (imagej.net/software/fiji).

#### Phagocytosis/trogocytosis of GUVs

50,000 macrophages were seeded in wells of an 8-well glassbottom plate (CellVis) in 100 *μ*L of RPMI 1640 medium. Post-seeding, cells were incubated at 37°C, 5 % *CO*_2_ for 3-4 hours before target addition. 100 *μ*L of low-tension GUVs or high-tension GUVs ( 1 million GUVs counted with an impedance-based cell counter (Scepter, SigmaAldrich)) were prepared with 4 *μ*M AlexaFluor647 anti-biotin IgG in PBS and allowed to incubate with gentle rotation for > 10 minutes. After washing, GUVs were added to macrophage-seeded wells on the stage top incubator of the microscope.

#### Time-lapse confocal microscopy of antibodydependent cellular phagocytosis

J774A.1 macrophages and Raji B lymphocytes were obtained from DSMZ (ACC170, ACC319). J774A.1 that stably express Lifeact-mScarlet3 [54] were cultured at 37°C with 5 % *CO*_2_ in DMEM (Wisent, 319-005) supplemented with 10 % of heat-inactivated FBS (Wisent, 090-150). Raji were cultured were cultured at 37°C with 5 % *CO*_2_ in RPMI-1640 (Wisent, 350-007) supplemented with 10 % of heat-inactivated FBS (Wisent, 090-150).24 hours before imaging, 100,000 macrophages were seeded into 18 mm circular #1.5 coverslips. Prior to the experiment, Raji cells were stained with Calcein AM viability dye (eBioscience, 65-0853-78) at 1 *μ*M for 30 minutes. Macrophages were transferred to a Chamlide CMB imaging chamber (Live Cell Instrument) and incubated in DMEM without phenol red, containing 25 mM HEPES (Wisent, 319-066). 200,000 Raji cells were added to the macrophages, and 2 *μ*g/ml of anti-human CD20 antibody (BioXcell, Rituximab) and 10 *μ*g/ml of anti-human CD47 antibody (BioXcell, B6.H12) were supplemented. Images were acquired every 20 seconds for 2 hours using a 40x 1.3 NA oil immersion objective on a Nikon Eclipse Ti2-E, equipped with a Yokogawa SCU-W1 spinning disk, a Hamamatsu Orca-Fusion BT sCMOS camera, and a stage-top incubator (Tokai Hit) to maintain cells at 37°C. Images were acquired using Nikon NIS Elements and analyzed using Fiji is just ImageJ 1.54p.

#### Lipid coating of hydrogel beads functionalized with TDA

First, unilamellar vesicles (LUVs) were prepared by the extrusion method. POPS (1-palmitoyl-2-oleoyl-sn-glycero-3-phospho-L-serine, Avanti Polar Lipids), POPC (1-palmitoyl-2-oleoyl-sn-glycero-3-phosphocholine, Avanti Polar Lipids) and Texas RedTM DHPE (Texas RedTM 1,2-Dihexadecanoyl-sn-Glycero-3-Phospho-ethanolamine, Invitrogen) dissolved in chloroform were mixed in a 80:19:1 molar ratio and dried. The dried lipid cake was then hydrated in PBS to a final concentration of 1 mM lipid suspension followed by 5 freeze-thaw cycles. Then, LUVs were prepared by 11-fold extrusion through a 100-nm polycarbonate filter (Avanti Polar Lipids) using a Mini-Extruder (Avanti Polar Lipids). Next, 50 *μ*L of vesicle solution were incubated for 45 min with 5 *μ*L of functionalized hydrogel beads followed 5 freeze-thaw cycles. Unbound vesicles were washed 5 times with PBS by centrifuging the solution for 1 min at 5000 g. Coated hydrogel beads were stored at 4ºC and used up to a week.

#### Zebrafish handling and maintenance

Zebrafish were maintained at the aquatic facility of the Parc de Recerca Biomèdica de Barcelona (PRBB) following the standard procedures proved by the Institutional Animal Care and Use Ethic Committee (PRBB–IACUEC) according to national and European regulations. All the experiments were performed following the principles of the 3Rs. Eggs were kept in E3 medium (5 mM NaCl,0.17 mM KCl, 0.33 mM CaCl2, 0.33 mM MgSO4) at 28°C. Embryos were staged based on morphological criteria and hours post fertilization (hpf). All embryos used in this study were obtained from the Tg(actb1:Lifeact-GFP) [55] zebrafish line.

#### Transplantation of synthetic targets

Acceptor embryos were dechorionated at sphere stage (4 hpf) and placed into a custom-made transplantation agarose mold (Adaptative Science Tools) containing Danieau’s solution (58 mM NaCl, 0.7 mM KCl, 0.4 mM MgSO4, 0.6 mM Ca(NO3)2, 5 mM HEPES). Synthetic targets were transplanted into the animal cap of acceptor embryos, close to the embryonic epithelial layer, using a Celltram Vario device (Eppendorf) in combination with an ES Blastocyst-pipette needle (Biomedical Instruments, VESbv-20-0-0-55).

#### Apoptosis induction of a subpopulation of cells

Apoptosis was induced in a subpopulation of embryonic cells as described previously [39]. In short, 2 pg of zebrafish bax mRNA [56] were co-injected with 100 pg of the membrane marker lyn-tdTomato mRNA into 16-32 cell stage embryos for a mosaic expression. mRNAs were synthetized from pCS2+ plasmids using the SP6 mMessage mMachine kit (Ambion, AM1340).

For more details of experimental and simulation method, see SI section S14.

## Supporting information

Suppementary Information

## II. CODE AVAILABILITY

The MATLAB code for analyzing confocal images and deriving particle shape is publicly available on https://gitlab.com/dvorselen/DAAMparticle Shape Analysis. The Python code for traction force analysis is available on https://gitlab.com/micronano-public/ShElastic.

## ACKNOWLEDGMENTS

N.S.G., incumbent of the Lee and William Abramowitz Professorial Chair of Biophysics, acknowledges support from the Israel Science Foundation (Grant No. 207/22) and the Harold Perlman Family. D.A.F. was supported by the National Science Foundation Center for Cellular Construction (DBI-1548297), the National Institutes of Health (R01GM134137), and the Chan Zuckerberg Biohub Investigator program. C.E.C. was supported by the James S. McDonnell Foundation Postdoctoral Fellowship. A.I. and S.P. acknowledge support from the Slovenian Research Agency (ARIS; project J3-60063 and programme P2-0232), the EU Horizon 2020 Marie Skłodowska-Curie Staff Exchange project *FarmEVs* (Grant No. 101131175), and COST Action CA22153. V.R. was supported by the Ministerio de Ciencia e Innovación (PID2020-117011GB-I00), HFSP (RGY0079/2020), the European Union’s Horizon Europe programme (BREAKDANCE, Grant No. 101072123), and the Austrian Science Fund (FWF; 10.55776/PIN3977225). M.B. acknowledges support from the Ministerio de Ciencia, Innovación y Universidades and Fondo Social Europeo (FSE; PRE2020-092691).

## Author contributions

SS and NSG designed the study and computational model. SS developed the code and performed simulations and data analysis. CEC, DAF, MKS, VJ, YP, DV, VR, and MB performed the experimental work, with VR and MB also generating lipid-coated hydrogel beads. AI and SP developed the original single-vesicle code. SS and NSG wrote the manuscript, and all authors reviewed and edited it.

