## Supplementary material for "From biting to engulfment: Target mechanics determines modes of phagocytosis through curvature–actin coupling": Suppementary Information

[1]*Department of Chemical and Biological Physics, Weizmann Institute of Science, Rehovot, Israel*
[2]*Department of Bioengineering, University of California Berkeley; Berkeley, CA USA*
[3]*Department of Molecular Biology and Biochemistry, Simon Fraser University, Burnaby BC, Canada*
[4]*Center for Genomic Regulation (CRG), The Barcelona Institute of Science and Technology, Barcelona, Spain*
[5]*Department of Cell Biology and Immunology, Wageningen University and Research, Wageningen, the Netherlands*
[6]*Laboratory of Physics, Faculty of Electrical Engineering, University of Ljubljana, Ljubljana, Slovenia*
[7]*University of California Berkeley/University of California San Francisco Graduate Group in Bioengineering, CA USA*
[8]*Division of Biological Systems and Engineering, Lawrence Berkeley National Laboratory; Berkeley CA USA*
[9]*Chan Zuckerberg Biohub; San Francisco CA USA*
[10]*Centre for Genomic Regulation (CRG), The Barcelona Institute of Science and Technology, 08003 Barcelona, Spain*
[11]*University of Innsbruck, 6020 Innsbruck, Austria*
[12]*Department of Physiology, Development and Neuroscience, Downing Site, University of Cambridge, Cambridge, UK*

(Dated: July 9, 2026)

### S1. THEORETICAL MODEL

We modelled the cell membrane as a three-dimensional vesicle which is described by a closed surface with $N$ vertices connected to its neighbours by bonds and it forms a dynamically triangulated, self-avoiding network, with the topology of sphere [8, 9]. $\boldsymbol{r_i}$ is the position vector of the $i$th vertex. All the lengths are measured in a scale of $l_{\min}$. There is a percentage $\rho = 100N_c/N$ of vertex sites that represent the curved membrane protein complexes (CMC), that induce cytoskeletal active forces. The vesicle energy has four components: The bending energy is given by,

$$W_b = \frac{\kappa}{2}\int_A (C_1 + C_2 - C_0)^2 \, dA, \tag{S1}$$

where, $\kappa$ is the bending rigidity, $C_1$, $C_2$ are the principal curvatures and $C_0$ is the spontaneous curvature. We consider the spontaneous curvature $C_0 = 1l_{\min}^{-1}$ for the CMC sites, represented in red and blue represents the bare membrane for which $C_0 = 0$. The protein-protein interaction energy is given by,

$$W_d = -w\sum_{i<j}\mathcal{H}(r_0 - r_{ij}) \tag{S2}$$

where $\mathcal{H}$ is the Heaviside function, $r_{ij} = |\boldsymbol{r_i} - \boldsymbol{r_j}|$ is the distance between protein sites, $r_0$ is the range interaction and $w$ is the strength. We set the parameter $w = 1k_BT$ throughout the paper. The energy due to the active force is given by,

$$W_F = -F\sum_i \hat{n}_i \cdot \boldsymbol{r_i} \tag{S3}$$

where $F$ is the magnitude of the active force, $\hat{n}_i$ is the outward unit normal vector of the $i$th protein site vertex and $\boldsymbol{r_i}$ is the position vector of the protein.

Finally, the adhesion energy due to an adhesive rigid substrate is given by,

$$W_A = -\sum_i E_{ad} \tag{S4}$$

where $E_{ad}$ is the adhesion strength, and the sum runs over all the adhered vertices to the substrate i.e., the vertices that are within a range of $1l_{min}$.

*
†

### S2. DETECTION OF OTHER VESICLES

We always started the vesicles are well separated in space, i.e., not intersecting each other. In multi-vesicle simulation, it is important to make sure that one vertex of a vesicle is not going into the other vesicles during the vertex move. If we pick a vertex from a vesicle that is very close to another vertex that belongs to other vesicle, then we check whether the vesicle is cutting the other vesicle. We need to check any such pair of vertices from different vesicles. Let the vertex of interest $\mathcal{V}_1$ (with which we want to make a Monte-Carlo (MC) vertex movement step), is very close to another vertex $\mathcal{V}_2$, then we exploit the information of tristar associated with the vertex $\mathcal{V}_2$. We have $T_2$ as the "tristar" (a list of triangles, see Fig. S1) associated with $\mathcal{V}_2$. The $i$th triangle of the tristar $T_2$ is presented by $T_{2i}$. Let $\overrightarrow{r_1}$ and $\overrightarrow{r_2}$ be the position vectors of $\mathcal{V}_1$ and $\mathcal{V}_2$ respectively. We calculate the projection of the relative position of $\mathcal{V}_1$ with respect to $\mathcal{V}_2$ that is $\overrightarrow{r_{12}} = \overrightarrow{r_1} - \overrightarrow{r_2}$ on the normals $\hat{n}_i$ of each triangles $T_{2i}$ as follows:

$$p_i = \overrightarrow{r}_{12} \cdot \hat{n}_i = (\overrightarrow{r_1} - \overrightarrow{r_2}) \cdot \hat{n}_i \tag{S5}$$

where $i$ runs through all the triangles in the tristar list $T_2$ of vertex $\mathcal{V}_2$. If any $p_i$ becomes negative, it means the vertex $\mathcal{V}_1$ intersected the triangle $T_{2i}$. Hence, we abort that vertex movement.

### S3. VESICLE-VESICLE INTERACTION

Two vesicles can interact with each other through adhesion and impart the active force due to the actin-cytoskeleton.

**Vesicle-vesicle adhesion**—Two different vesicles can adhere to each other. Let $E_{\rm ad}$ be the adhesive energy per node for cell-cell adhesion. The interaction between two vesicles through adhesion becomes effective only when the two vesicles come very close and within a finite range of adhesion $1l_{\rm min}$. Therefore, if a vertex of vesicle 1 comes within the range of cell-cell adhesion of any of the vertices belonging to another vesicle (say vesicle 2), then we count $E_{\rm ad}$ as the cell-cell adhesion energy in the energy calculation of the vertex of interest that belongs to the vesicle 1 (See Fig.1D).

**Force applied on the other vesicle**— A vesicle can impart active force on the other vesicle. If a vertex is within the range of interaction (The interaction range is set to 1 $l_{\rm min}$) of other protein vertices belonging to the other vesicle, a vectorially added effective force is applied on the former vertex. If a set $\mathcal{S}$ of vertices belongs to another vesicle are within the interaction range of a vertex of interest $\mathcal{V}_i$ then the force applied on the vertex $\mathcal{V}_i$ is given by,

$$\overrightarrow{F}^{\rm int} = \sum_{V_i \in \mathcal{S}} \overrightarrow{F}_{V_i}. \tag{S6}$$

The interaction between the vesicles is mediated by the active forces applied by the others vesicles. Any vertex feels the active force due to the other vesicle's active protein sites. The effect of interaction force is included by adding the energy cost due to the interaction force given by,

$$W_{\rm int} = \overrightarrow{F}^{\rm int} \cdot \overrightarrow{dr}. \tag{S7}$$

where, $\overrightarrow{dr}$ is the monte carlo movement of the vertex of interest.

### S4. CALCULATION OF ADHERED AREA FRACTION OF THE TARGET VESICLE

To calculate the adhered area fraction of the target vesicle, we used a simple method by counting the number of nodes that are in the vicinity of the nodes of other vesicles and dividing it by the total number of nodes. We calculate it for different cases as follows:

**Adhesive area fraction calculation for the triangulated vesicle**— To calculate the adhesion area fraction, we scan through all the vertices of a vesicle to check if the vertex is within the adhesion range from any other vesicle or the adhesive substrate. If the vertex is within the adhesion range to any other adhesive surface, then we count it as an adhered vertex. Finally, we find the approximate adhesive area fraction by dividing the number of adhered vertices $N_{\rm ad}$ by the total number of vertices $N$.

**Adhesion fraction calculation for rigid sphere**— We do not have the information for the vertices for the case of a rigid sphere. To make a similar calculation to the calculation used in the vesicle with a triangulated membrane, we generated 847 vertex points (same as the number of vertices on the target vesicle) on the sphere using a Fibonacci lattice method.

Let's say, we shall place $N^T$ number of vertices on a sphere of radius $R$, centred at $(x_0,\ y_0,\ z_0)$ using the Fibonacci lattice. We shall generate $N^T$ points given by,

$$x_i = x_0 + R\left(1 - \frac{2(i-1)}{(N^T-1)}\right)$$
$$y_i = y_0 + \sqrt{(R^2 - x_i^2)}\sin\phi$$
$$z_i = z_0 + \sqrt{(R^2 - x_i^2)}\cos\phi \tag{S8}$$

where, $\phi = \pi(3-\sqrt{5})$ is the golden angle and the integer index $i$ runs from 1 to $N^T$. We placed 847 points on a sphere of radius 10 $l_{\text{min}}$. Then, we do the same calculation as before.

### S5. VERIFICATION OF RADIUS OF CURVATURE NEAR THE CONTACT WITH A SINGLE VESICLE

A single vesicle is allowed to spread on a flat adhesive substrate (Fig. S2A). After getting to the steady state, we calculated the radius of curvature near the contact when the area and the volume of the vesicle are conserved. It can be theoretically estimated from the energy terms if there are no curved proteins and no active forces. Theoretically, the radius of curvature is given by $R = \sqrt{\kappa/2E_{\text{ad}}}$ [4]. We analyzed a two-dimensional cross-section of the vesicle shape, let's say $x-z$ plane as shown in Fig. S2A. Then, we approximately identified the contact region and then fit a circle of radius $R$ that gives the radius of curvature near the contact point, as shown in Fig. S2B. Since the numerical results may fluctuate, we have found the value of $R$ by taking an average of the fit after rotating the vesicle about the $z$ axis as it has rotational symmetry about the $z$ axis. We took ten realizations by rotating the vesicle in steps of $36^o$. We can estimate an error by taking the statistical error bar.

One can find the contact angle $\theta_c$ by fitting the straight line near the contact as shown in Fig. S2B. We did a similar angular averaging to calculate the contact angle.

A single vesicle is allowed to spread on an adhesive substrate for different adhesion energy $E_{\text{ad}}$ parameters and different CMC densities $\rho = 0,\ 3.46\%,\ 6.93\%$. The shapes for all these different cases when the total volume of the vesicle is conserved are shown in Fig. S3A. We showed the case when the vesicle volume is not conserved in Fig. S3B. We have shown the steady-state adhered area fraction for all these cases in Fig. S3C. There is a higher adhered area fraction when the adhesive energy $E_{\text{ad}}$ per node is higher. We also see that the curved-protein percentage is driving the spreading, by reducing the bending energy cost near the contact line. When the volume is not conserved, we find that the vesicle can lose its volume to assume a sheet-like shape and spread more, as shown in Fig. S3C.

We have shown the two-dimensional side cut of the vesicle on $x-z$ plane and the fitted circle in order to calculate the radius of curvature in lime (See Fig. S3D). We verified the radius of curvature $R$ for $E_{\text{ad}} = 1k_BT$ and $2k_BT$ when volume is conserved and the curved proteins are absent as shown in Fig. S3E. The theoretical estimation is given by the grey solid line .

### S6. VERIFICATION OF RADIUS OF CURVATURE NEAR THE CONTACT WITH TWO IDENTICAL VESICLES

Next, we consider two identical vesicles that adhere to each other. Again, we find the radius of curvature near the contact and validated with the previous work [4]. Though the technique to calculate the radius of curvature is similar to the case of the single vesicle, you need to orient the vesicles properly before making any observation. We first draw a straight line joining the center of mass of the two vesicles, as shown in Fig. S2C. We then align this line to the $z$ axis by rotation. We calculated the radius of curvature near the contact for the cases of no CMC and with volume conservation, for two different parameters for the adhesion energy $E_{\text{ad}}$ per node. The snapshots for two vesicles spreading on each other are shown in Fig. S4A. The adhesive area fraction is increased as the adhesive energy per node is increased (See Fig. S4B). Also, the adhesion area fraction is larger in the presence of the CMC. The volume conservation hinders the spreading, with the adhered area fraction increasing with increasing CMC concentration and increasing adhesive energy $E_{\text{ad}}$ per node. Fig. S4C shows the side cut of the two vesicles adhering to each other for the no CMC and volume conservation case. The verification of the radius of curvature near the contact point is shown in Fig. S4D.

### S7. ADHESION DYNAMICS BETWEEN TWO VESICLES WITH PASSIVE CMC AND NO VOLUME CONSERVATION

In Fig.2 we show the steady-state final shapes of two identical vesicles adhered to each other for different strengths of adhesion energy parameter $E_{ad}$ and passive CMC. In Fig.S5 we plot the time evolution of the energy terms for the two vesicles during these processes. We show that above a critical adhesion energy of $\sim 1.2k_BT$ there is spontaneous breaking of the symmetry, with one of the vesicles experiencing a large increase in its bending energy (Fig.S5A), which is of course compensated by the large increase in the negative magnitude of the adhesion energy (Fig.2B,C). The vesicle that deforms into a cap-shape forms a ring cluster of its CMC, which indeed provides a larger interaction energy between the proteins (Fig.S5B).

### S8. BENCHMARK OF PHAGOCYTOSIS WITH A RIGID SPHERICAL PARTICLE

Next, we have two non-identical vesicles and let them interact through adhesion, and the active forces that mimic the actin cytoskeleton. One vesicle is bigger in size and contains CMC, representing a cell-like vesicle. The other vesicle is smaller in size and has no CMC for simplicity, serving as the target-vesicle. We kept the bending rigidity of the cell-like vesicle $\kappa = 20k_BT$ throughout the paper. Therefore, whenever we talk about the bending rigidity $\kappa$ in the context of the phagocytosis process, it is the bending rigidity of the target vesicle.

Before we explore the complex behaviours exhibited by the very soft objects due to their big deformations, we validate the model by comparing two cases: We consider a completely rigid spherical object as a target [7], and compare the dynamics to the case of a very rigid target (high bending rigidity $\kappa = 2000k_BT$) vesicle of the same radius. We simulate these two cases with a model cell-like vesicle containing wither passive ($F = 0k_BT\ l_{\mathrm{min}}^{-1}$) or active ($F = 2k_BT\ l_{\mathrm{min}}^{-1}$) CMC.

We find the adhered area fraction is very similar for the completely rigid object and the target vesicle with very high bending rigidity $\kappa = 2000k_BT$ for both the passive and active cases, respectively, as shown in Fig. S6E-F. We have also found that the active vesicle is more efficient in complete engulfment compared to the passive vesicle, as shown in Fig. S6G. We have shown the different stages of the engulfment processes with snapshots of the interaction between two vesicles in Fig. S6A-D for four different cases.

### S9. VESICLE AREA DEFORMATION AND FLUCTUATIONS DURING PASSIVE ENGULFMENT

In Fig.3 we show the engulfment of vesicles of different bending rigidities by cell-like vesicle that contains passive CMC. The engulfment involves deformations of the target vesicle, which are quantified as follows: First, we find the centroid $\overrightarrow{r}_0$ of the vesicle by averaging the position vectors of all the vertices. Next, we find the distances of all the vertices from the centroid that is given by,

$$\tilde{r_i} = |\overrightarrow{r}_i - \overrightarrow{r}_0|. \tag{S9}$$

Next, we find the relative standard deviation of these distances $r_i$ is given by,

$$\Delta R/R = \sqrt{\langle \tilde{r_i}^2 \rangle - \langle \tilde{r_i} \rangle^2} / \langle \tilde{r_i} \rangle. \tag{S10}$$

A sphere has a non-roundedness measure of zero, and it increases as the vesicle becomes more non-spherical.

The non-roundness parameter for the target vesicle is shown in Fig.3C. Similarly, changes to the vesicle's area during this engulfment process are shown in Fig.S7.

### S10. DETACHMENT OF THE CELL-LIKE VESICLE FROM THE TARGET AFTER A PUSHING EVENT

In Fig. 4D we showed that for intermediate target vesicle rigidities the cell-like vesicle ends up pushing it. This process ends with the cell-like vesicle detaching from the target vesicle and retracting the long protrusion that formed. In Fig.S8 we plot the bending energies of the two vesicles during this process, and the sum of the adhesion and protein-protein interaction energy for the cell-like vesicle. We see that as the pushing event progresses, there is an increase in the bending energy associated with the long protrusion of the cell-like vesicle ($E_{b1}$ in Fig.S8), while the adhesion and protein-protein binding energies which compensate for it decrease. This increasing energy cost eventually drives the detachment, and retraction, which decreases the bending energy cost.

### S11. EFFECT OF HOLDING THE TARGET ON THE ENGULFMENT PROCESS

We have found that in the regime of intermediate rigidity, the cell-like vesicle pushes the target vesicle and ultimately leaves it without biting or engulfing it (Fig. 6E, Fig. 4D). Here we simulated the dynamics when an external constraint confines the target vesicle. For example, in an "in vitro" experiment a target may be held by a pipette [1], or cells may be confined by surrounding cells inside tissues.

To mimic such a confinement effect, we designated a patch of vertices on the target vesicle that are not allowed to move. We choose a patch of vertices centered at the pole of the target vesicle away from the cell-like vesicle (black patch on the right-most edge of the target vesicle in Fig. 6F). This is a patch of depth of $l_{\min}$ from the pole of the target vesicle. We found that this confinement can change the dynamics from pushing to engulfment or biting (trogocytosis), as shown in Fig 6 (E-F) and see the Movie S11.

We showed how different quantities, such as the adhered area fraction, deformation, tangential force fraction and normal force fraction evolve over MC steps as shown in Fig. 6 (G-I). Holding a patch of the target vesicle changes the entire engulfment dynamics by directing more of the active force to be tangential to the target vesicle, therefore much more efficient in driving engulfment.

### S12. EFFECT OF INTERNAL PRESSURE OF THE TARGET CELL

When an osmotic pressure difference $p$ exists between the inside and the outside of the target vesicle, an additional energy cost term is included [2]

$$\Delta E_{\mathrm{p}} = p dV \tag{S11}$$

Here, $dV$ is the change in volume due to the Monte Carlo moves (e.g. vertex movement, bond flip). The positive internal pressure $p$ creates more tension on the membrane, and the vesicle gets inflated.

We have systematically studied the effect of the internal osmotic pressure of the target vesicle on the engulfment process, while keeping the bending rigidity of the target-like vesicle at $\kappa = 20k_BT$. We found three dynamical phases (Fig. S9), identical to the phases found for varying the bending rigidity (Fig.4). As the internal pressure increases, the target vesicle membrane is under higher tension, which opposes the deformation and keeps it more spherical. Increasing internal osmotic pressure in the target vesicle therefore affects the engulfment process in an identical way to increasing the bending rigidity of the target vesicle (Fig.4).

We next calculated a measure of the membrane tension due to the internal osmotic pressure $p$, as shown in Fig.S10. As both the volume and area of the vesicle are changing, we introduced the volume-normalized area $\tilde{A}$ given by,

$$\tilde{A} = \frac{A}{V^{2/3}} \tag{S12}$$

where, $A$ and $V$ are the area and the volume of the vesicle. The time evolution of $\tilde{A}$ is shown in Fig. S10A for the cases of internal pressures of $p = 0.1,\ 0.5$ and 10, during the interaction of the target with an active cell-like vesicle (as in Fig.7G-I). Next, we calculated the probability distribution of the volume-normalized area $\rho(\tilde{A})$ (Fig.S10B). As pressure increases, the distribution function has a very narrow and sharp peak. A sharper peak implies a lower standard deviation $\sigma_{\tilde{A}}$ of the area fluctuations for higher internal pressure $p$ as shown in Fig. S10C. Lower area fluctuations are equivalent to higher membrane tension.

### S13. IMPLEMENTATION OF BULK MODULUS

In order to compare with the experiments of phagocytosis of elastic beads [10, 13] at a high resolution (Fig. S12), we need to implement a description of the bulk elasticity of the target vesicle.

We implemented a bulk modulus for the target cell in a simple way. The desired shape for the vesicle is a sphere with a radius $r_0 = 10\ l_{\min}$. All the vertices feel a force or cost some energy if it deviates from that shape. The force is given by,

$$\mathbf{F}_i^{\mathrm{bulk}} = \kappa_{\mathrm{bulk}}(r_i - r_0)\hat{n}_{\mathrm{bulk}} \tag{S13}$$

where, $r_i$ is the distance of the center of mass from the $i$th vertex, and $\hat{n}_{\mathrm{bulk}}$ is the unit vector directed to the center of mass from the $i$th vertex. We added the energy due to the bulk modulus of the target that is set to $\kappa_{bulk} = 0.5\ k_BT/l_{\min}^2$. We simulated the engulfment of soft target of $\kappa = 20k_BT$ by a cell-like vesicle and measured

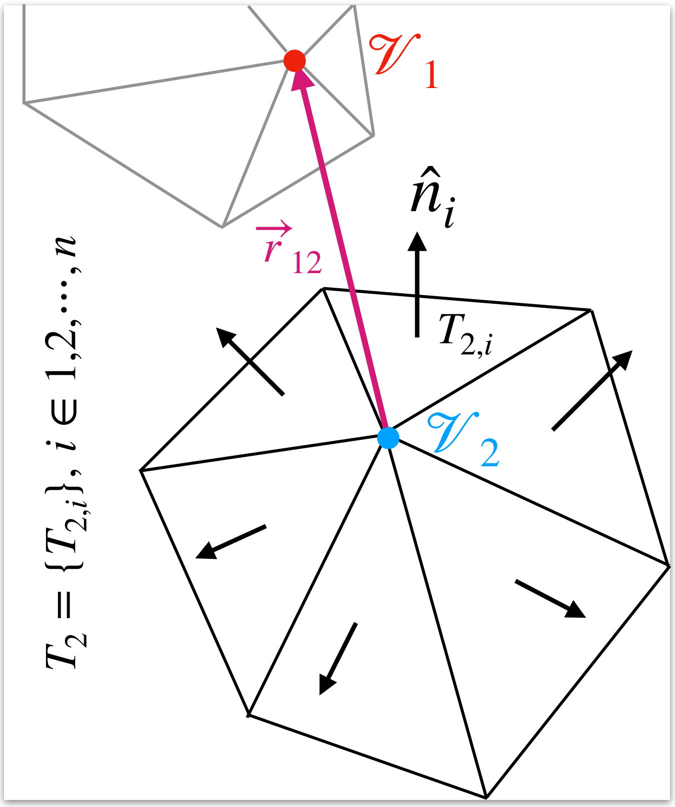


FIG. S1. Demonstration of the detection of an overlap between two vesicles. Two vertices $\mathcal{V}_1$ and $\mathcal{V}_2$ belong to two different vesicles separated by the position vector $\vec{r}_{12}$. We check a MC change to the position of the vertex $\mathcal{V}_1$, given that the vector $\vec{r}_{12}$ magnitude is less than the minimum length scale $l_{\min}$. Keeping the vertex $\mathcal{V}_2$ at the center, the neighbouring triangles create a triangle list denoted by the set $T_2$. We check the dot product of the normals of the member triangles with the position vector $\vec{r}_{12}$. If any of the dot products is negative, then the two vesicles are overlapping.

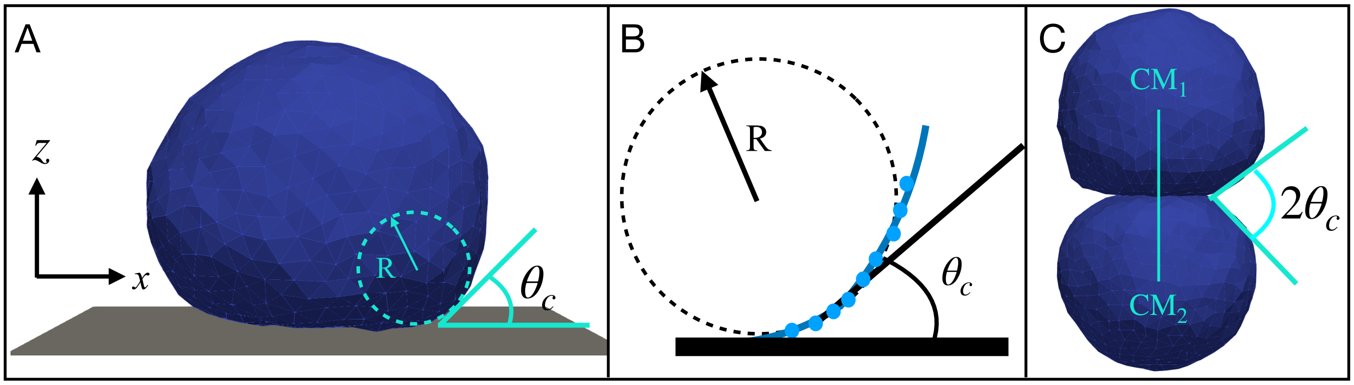


FIG. S2. (A) A vesicle is adhered on an adhesive substrate. We find the cross section of the vesicle on the $x - z$ plane. Near the adhesion region we find the radius of curvature $R$. We find the angle of contact by making a linear fit with the points near the contact line: (B) Here, the relevant points along the vesicle surface are shown in blue dots. They are used to make a circular fit to find $R$, and the linear fit to find the contact angle $\theta_c$. (C) For two vesicles adhering to each other, first we join their centers of mass ($CM_1$ and $CM_2$) and align it along $z$ direction. Then we applied the same procedure to find $R$ and $\theta_c$ as in (B). As the system has rotational symmetry around the $z$-direction, we averaged over 10 cases by rotating the system in steps of $36^o$ around the $z$ direction.

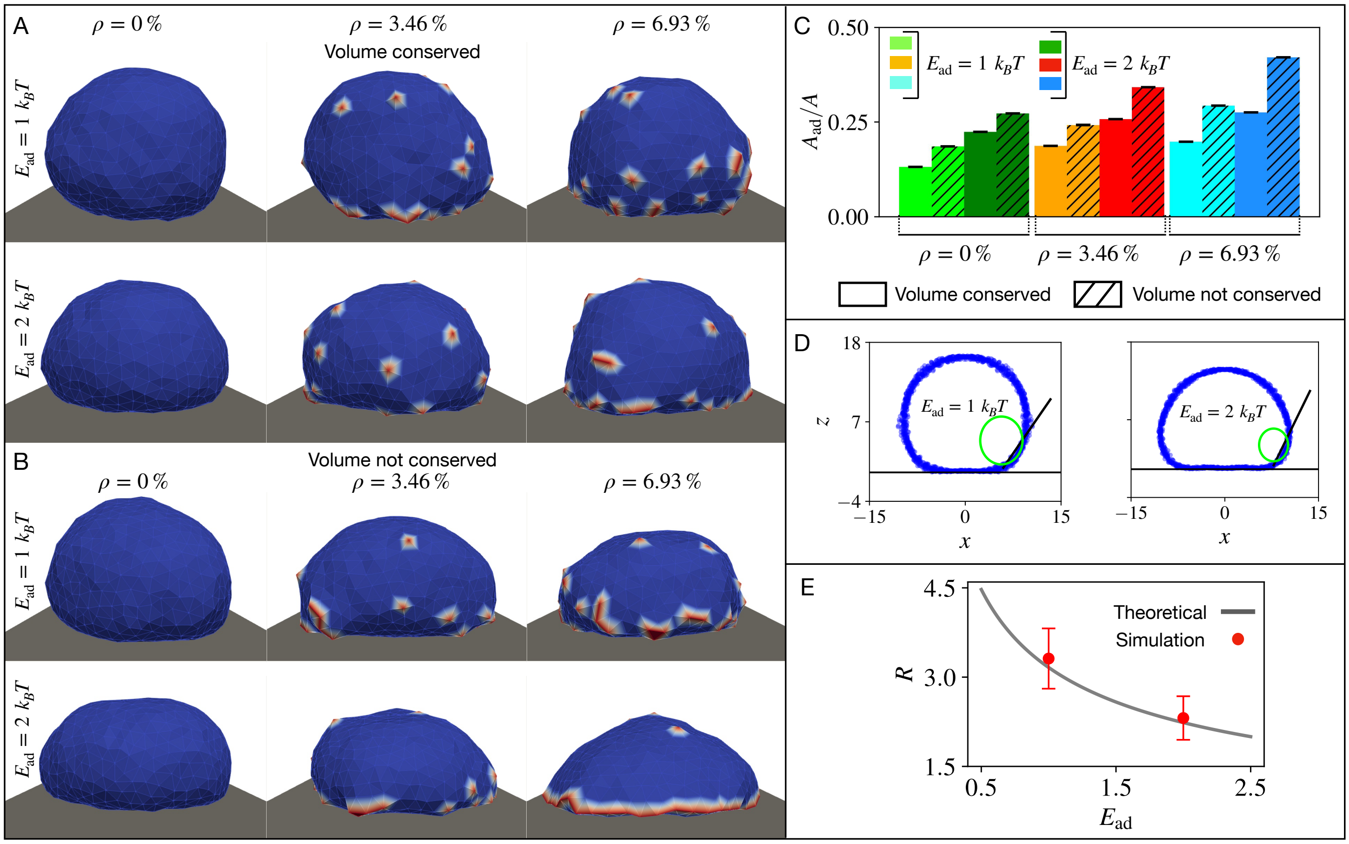


FIG. S3. Verification of the radius of curvature near the contact line of a vesicle on the flat substrate. A) Snapshots of the vesicle adhering to the flat substrate when the volume and the area are conserved, for different concentrations of CMC (passive) and adhesion energies. B) Same as (A) when the volume conservation constraint is removed. C) Adhered area fraction is shown as a bar plot for different cases shown in (A,B). D) Numerical extraction of the radius of curvature near the substrate-vesicle contact line from the simulated shapes. E) Verification of the predicted dependence of the radius of curvature (as defined in Fig.S2) on the adhesion energy [4] for two cases of $E_{\text{ad}} = 1k_BT,\ 2k_BT$, when there are no CMC ($\rho = 0\%$) and the volume is conserved.

the actin strength, radial deviation, and the normal force along the formed phagocytic cup or actin ring. We compared these quantities with the experimental data which is done with artificial DAAM particles, as shown in Fig. S12 (A-B). We compared the total normal force with the fraction of the adhesion area of the target or the fraction of engulfment. The overall agreement is very good. Note that the actin ring in the experiments is necessarily wider than the ring of CMC in the model, since the CMC denote only the leading edge nucleators of actin and not the whole actin network that forms behind them.

In the experiments at the bead's pole pointing at the engulfing cell a pulling force acting on the bead is measured (Fig. S12A), which may arise from contractile forces pulling on these adhesion sites, or due to the effect of actin treadmilling emanating from the leading edge, exerting forces that pinch the bead at this pole (similar to actin treadmilling-induced forces that are involved with endocytosis [6]). This effect is absent from our simple model.

### S14. DETAILS OF EXPERIMENTAL AND SIMULATION METHODS

**Lipids**—Phosphocholine (PC) lipids (Avanti Polar Lipids) were used as purchased without further purification. Lipid stock solutions in chloroform contained a ternary mixture of 98 mol% POPC, 1 mol% biotin-PE, and 1 mol% PEG2K DSPE. GUVs are diluted in an ionic solution of PBS and all lipids in our mixtures are zwitterionic. We added PEG2K DSPE to block GUVs from aggregating in the charge-screened PBS solution.

**Antibodies**—Antibodies used to opsonize GUVs were purchased from Santa Cruz Biotechnology and used without further labeling or purification. Biotin was bound by AlexaFluor647-labeled anti-biotin mouse IgG (clone BK-1/39,

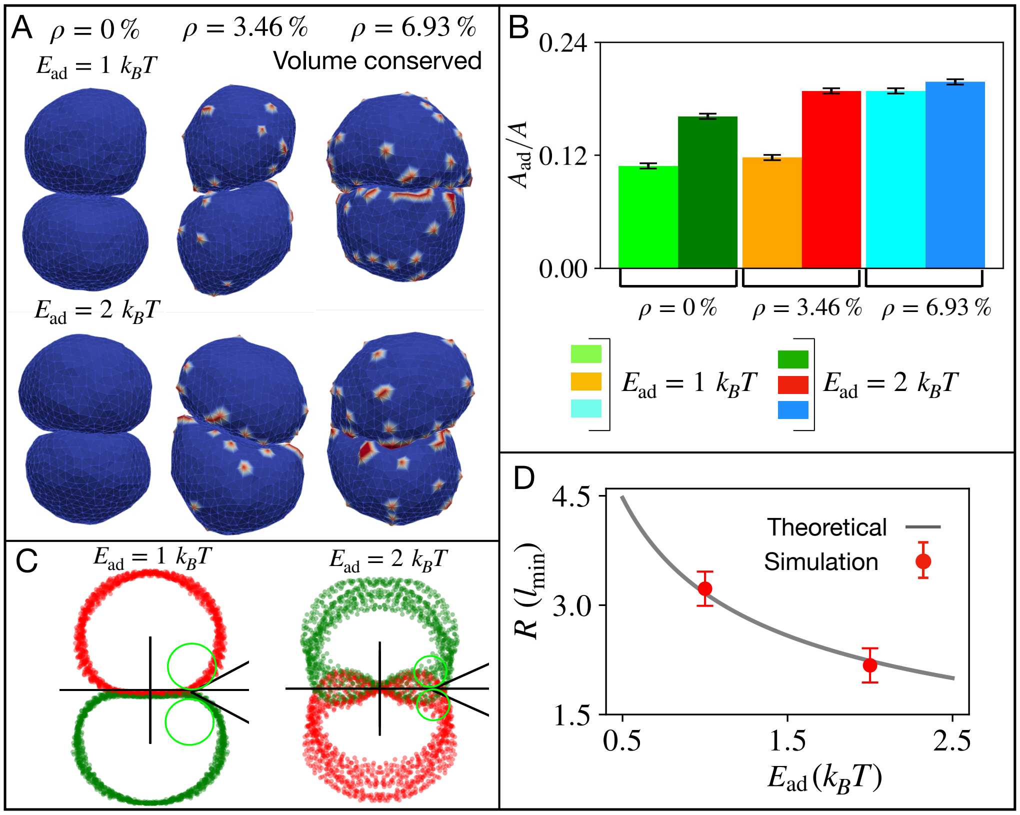


FIG. S4. Verification of radius of curvature near the contact line of two adhered symmetric vesicles when the volume and area are kept constant. A) Snapshots of two vesicles adhering to each other for three different CMC densities ($\rho = 0\%$, 3.46%, and 6.93%), and different adhesion energies $E_{\rm ad} = 1k_BT,\ 2k_BT$. B) The adhered area fraction is shown as a bar plot. C) The numerical calculation of the radius of curvature near the contact line between the two adhered vesicles for the case of $\rho = 0\%$. D) Comparison of the numerical estimates of the radius of curvature compared with the theoretical prediction $R_{\rm th} = \sqrt{\frac{\kappa}{2E{\rm ad}}}$ shown in grey line. We used the number of vertices $N = 722,\ \kappa = 20k_BT,\ w = 1k_BT,\ c_0 = 1l_{\rm min}^{-1}$ for both vesicles.

Santa Cruz Biotechnologies).

**RAW 264.7 cell culture**—RAW 264.7 murine male macrophage-like cell line was obtained from and authenticated by the UC Berkeley Cell Culture Facility. Cells were cultured in RPMI 1640 media (Corning) supplemented with 10 % heat-inactivated fetal bovine serum (HI-FBS, Thermo Fisher Scientific) and 1 % Pen-Strep (Thermo Fisher Scientific). RAWs were cultured in non-tissue culture-treated 10 cm dishes (VWR) at 37$^o$C, 5 % $CO_2$.

**Stable LifeAct GFP RAW 264.7 cell line**—HEK293T cells were grown in a 6-well plate to 80% confluency, and 160 ng VSV-G, 1.3 $\mu$g CMV 8.91, and 1.5 $\mu$g target pHR LifeAct GFP expression vector were transfected into HEK293T cells using TransIT-293T transfection reagent (Mirus Bio). Viral supernatants were collected 60 hours after transfection and spun at 4000 G to remove HEK293T cells. Viral supernatant was stored at 4$^o$C for no longer than 48 hours prior to infection. For lentiviral infection, 500 $\mu$L of viral supernatant was added to 5e5 RAW 264.7 macrophages along with 4 $\mu$g/mL polybrene, and cells were spun at 400G for 25 minutes at 37$^o$C and then resuspended and plated in a 6-well plate. Viral media was replaced with fresh growth media 24 h after infection. Cells were sorted via fluorescence-activated cell sorting on an Influx Cell Sorter (Beckton-Dickinson), and a population of cells expressing LifeAct GFP was expanded and frozen for later use.

**Synthesis of deformable acrylamide acrylic acid particles**—Hydrogel particles were synthesized as previously described. Briefly, acrylamide mixtures of acrylamide (AAm), acrylic acid (AAc), crosslinker $N,N'$-methylenebisacrylamide (BIS), 150 mM NaOH, 0.3% (v/v) tetramethylethylenediamine (TEMED), 150 mM MOPS (prepared from MOPS sodium salt, pH 7.4) were prepared. Total mass concentration of acrylic components ($C_{\rm AAm} + C_{\rm AAc} + C_{\rm BIS}$) was 100 mg/mL respectively 200 mg/mL and crosslinker concentration ($C_c = m_{\rm BIS}/(m_{\rm AAm} + m_{\rm AAc} + m_{\rm BIS})$) 0.64% respectively 5.06%, for 1.3 kPa and ~80 kPa particles. Prior to extrusion, the mixture was

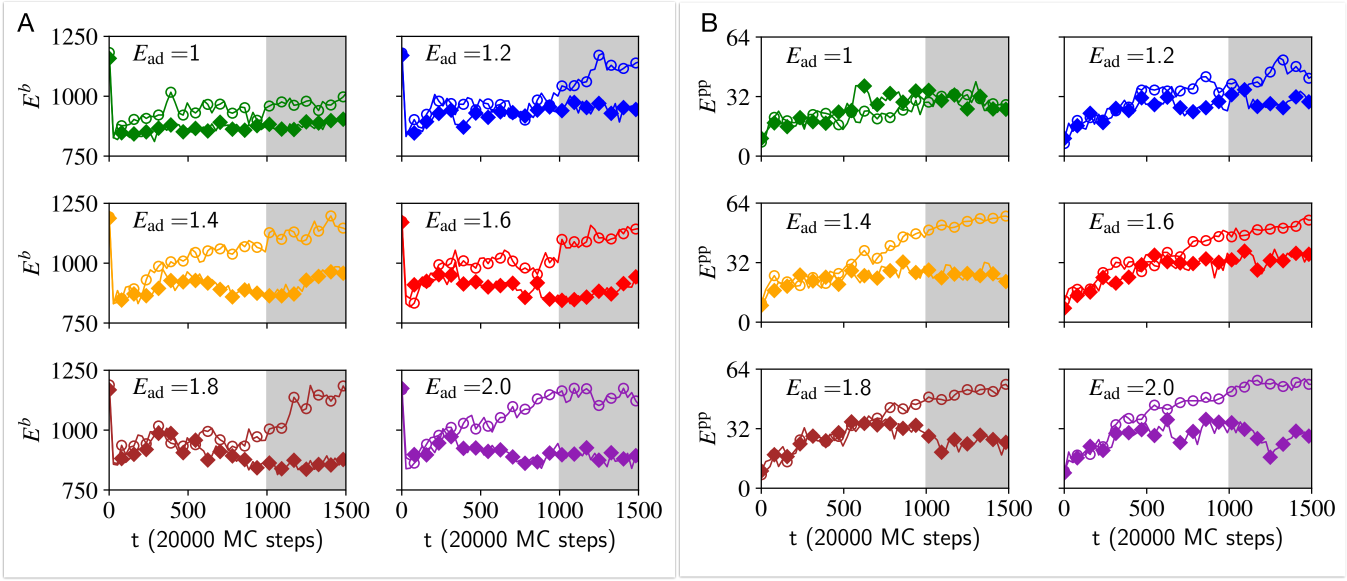


FIG. S5. Energy terms during the adhesion dynamics between two vesicles shown in Fig.2 (open symbols for the top vesicle 2, and filled symbols for the bottom vesicle 1). A) The asymmetry in the bending energies $E^b$ for two identical vesicles without volume conservation as the adhesive energy strength increases from $1k_BT$ to $2k_BT$. The bending rigidity for two vesicles is set to $\kappa = 20k_BT$ and the protein percentage is set to $\rho = 6.92\%$ for both vesicles. B) The asymmetry in protein-protein interaction energy $E^{\rm pp}$ for the same cases as in (A). Each vesicle consists of 722 vertices.

degassed for 15 min and kept under nitrogen atmosphere. Tubular hydrophobic Shirasu porous glass (SPG) membranes of 20 mm length with pore size (diameter) 1.1 $\mu$m respectively 2.4 $\mu$m were used for soft and stiff particles. Membranes were sonicated under vacuum in HPLC grade n-heptane to remove gas trapped in the membrane. The membranes were mounted on an internal pressure micro kit extruder (SPG Technology) and immersed into an oil phase (∼125 mL) consisting of hexanes (99% ACS reagent, mixed isomers) and 3% (v/v) Span 80 (Sigma Aldrich, S6760). 10 mL of gel mixture was extruded through SPG membranes under nitrogen pressure of ∼30 kPa, while the oil phase was continuously stirred and kept under nitrogen atmosphere in a 3-neck water-jacketed flask. After completion of extrusion, the emulsion temperature was increased to 60°C. Once the temperature equilibrated, DAAM-particle polymerization was induced by addition of ∼225 mg 2,2'-Azobisisobutyronitrile (AIBN) (1.5 mg/mL final concentration). The polymerization reaction was continued for 3 h at 60°C and then at 40°C overnight. Polymerized particles were subsequently washed (5× in hexanes, 1× in ethanol), dried under nitrogen flow for ∼30 min, and resuspended in PBS, pH 7.4. Soft and stiff particles had diameters of 8.9 respectively 8.2 mm, as determined by phase-contrast imaging and image analysis in ImageJ.

**Microparticle traction force microscopy (MP-TFM) analysis of phagocytosis of DAAM-particles—** Experimental data was obtained from prior conducted experiments [11] in which RAW 264.7 macrophages were exposed to IgG-functionalized 1.3 kPa deformable poly-Aam-co-AAc microparticles (DAAM-particles), stained for F-actin using Alexa Fluor-488 conjugated phalloidin, and imaged by confocal microscopy [12]. DAAM-particle 3D shape reconstructions and force analysis were performed as previously described [3, 14]. Briefly, the inverse problem of inferring the traction forces $\boldsymbol{T}$ is solved iteratively until a minimal gradient tolerance is reached. During this optimization, an ideal sphere with the same particle volume of the measured individual particle is subjected to a trial displacement field (u) to exactly match the surface of the experimentally observed shape of the DAAM-particle, while minimizing the cost function:

$$f(u) = E_{el} + \alpha R^2(T; \partial\Omega_t) + \beta E_{pen}(T) \tag{S14}$$

where $R(T; \partial\Omega t)$ represents the residual cellular forces exerted outside of the cell-target contact region, defined from the phalloidin and immunostaining. The elastic energy ($E_{el}$) penalizes unphysical solutions in which larger forces producing the same shape, while $\beta E_{pen}$ serves as an anti-aliasing term. The weighing parameters, a (residual traction) and $\beta$ (anti-aliasing), were both set to 1. Spherical harmonic coefficients up to $l_{max}$= 20 were utilized and normal forces were evaluated on a 21 × 41 grid.

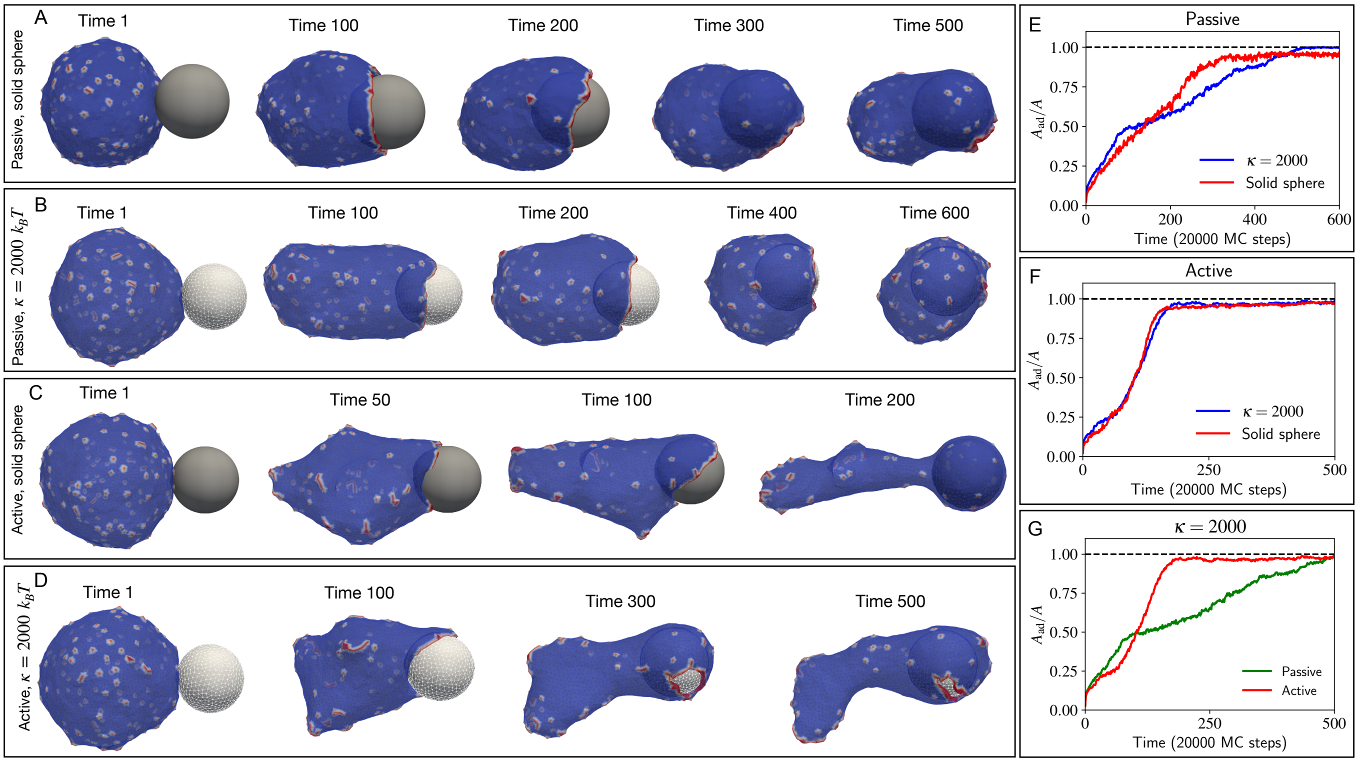


FIG. S6. The snapshots of the phagocytosis process when the big vesicle with passive proteins (no. of vertices $N = 3127$, no. of passive proteins $N_p = 150$, and bending rigidity $\kappa = 20\ k_BT$) tries to engulf a (A) rigid sphere of radius $R = 10\ l_{\min}$ and (B) a vesicle (nearly a sphere of radius of $R \approx 10\ l_{\min}$ in shape) with high bending rigidity $\kappa^T = 2000 k_BT$. Next, the big vesicle is active, it can exert force on the target. The active force parameter $F = 2\ k_BT\ l_{\min}^{-2}$. The snapshots are shown when the target is (C) a rigid sphere of radius $R = 10\ l_{\min}$ and (D) a vesicle with high bending rigidity $\kappa^T = 2000 k_BT$. Comparison of the adhesive area fraction of the target between the rigid sphere and the vesicle of bending rigidity $\kappa^T = 2000\ k_BT$ when the engulfing vesicle is (E) passive $F = 0$ and (F) active case $F = 2\ k_BT\ l_{\min}^{-2}$. (G) Comparison of the adhesive area fraction of the target between the passive and active engulfment.

**Interaction between the vesicles**—Two vesicles were left to interact through adhesion. The adhesive energy per node between two vesicles is set to $E_{\text{ad}} = 2k_BT$. If the vertices from different vesicles come within the interaction range (set to $l_{\min}$) then this adhesive energy is taken into account. The process is passive when the interaction is solely through the adhesion between them, and there is no active force imparted on the target vesicle by the bigger cell-like vesicle. An active force is imparted by a CMC node in the cell-like vesicle on a vertex in the target vesicle. To calculate the tangential and normal force imparted, we calculate the interaction force first. Let $V_i$ be the vertex of interest in the target vesicle, and $\hat{n}_i$ be its outward normal (computed by taking the average of the normals for all the triangles having the vertex in common). The normal and tangential components of the interaction force due to the sum of active forces acting on this vertex, $\mathbf{F_i}$, are given by,

$$\begin{aligned} \mathbf{F}_\perp &= (\mathbf{F_i} \cdot \hat{n}_i)\hat{n}_i, \\ \mathbf{F}_\parallel &= \mathbf{F_i} - (\mathbf{F_i} \cdot \hat{n}_i)\hat{n}_i. \end{aligned} \tag{S15}$$

**Setup for the engulfment simulation**— In all the simulations of the engulfment process, we construct two vesicles: i) a big cell-like vesicle with 3127 vertices, out of which 150 vertices represent the curved proteins with intrinsic mean curvature $c_0 = 1\ l_{\min}^{-1}$, the protein-protein interaction energy is set to $w = 1\ k_BT$, and ii) a smaller target vesicle made of 847 vertices with no curved proteins. At the initial time, the two vesicles were nearly spherical in shape and placed very near to each other, within the adhesion distance. The bending rigidity for the cell-like vesicle is set to 20 $k_BT$ throughout the paper. We varied the bending rigidity of the target vesicle from 20 $k_BT$ to a very high value of 2000 $k_BT$, which is nearly a rigid sphere.

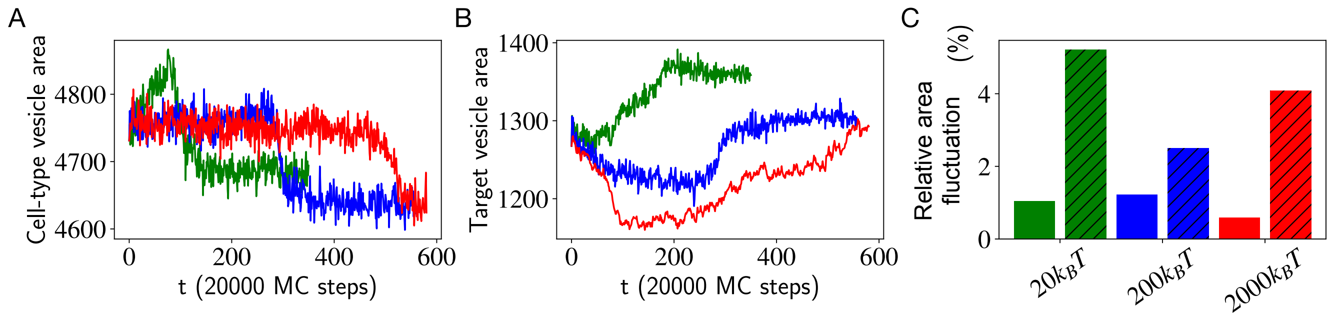


FIG. S7. The evolution of the vesicles' area during the engulfment simulations for the passive case (Fig.3): A) The cell-like vesicle area over time, for three different bending rigidity values of the target vesicle: $\kappa = 20,\ 200,\ 2000$ respectively in the units $k_BT$ (green, blue and red respectively). B) The target vesicle's area over time, as in (A). C) The relative fluctuation of the area for both vesicles is calculated with respect to the initial area of the vesicle. The smooth-color bars are for the cell-like vesicle, while the diagonal hatched bars are for the target vesicle.

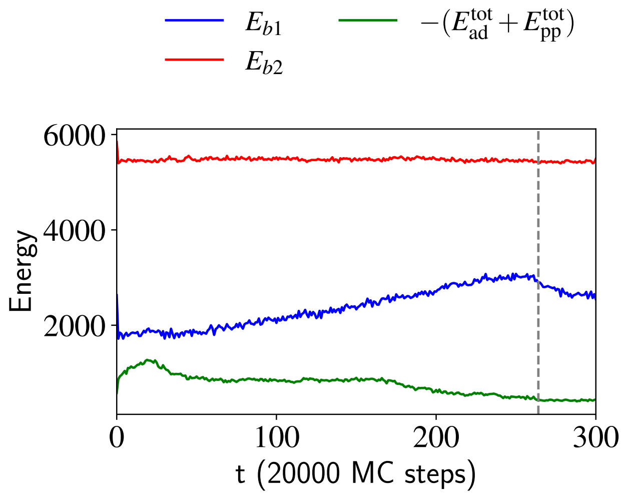


FIG. S8. The timeseries for the energy terms for the case of pushing (Fig.4D), when the bending rigidity of the target is $\kappa = 200\ k_BT$. The bending energy of the target vesicle is nearly constant over time (red). We show the competition between the bending energy of the cell-like vesicle (blue) and its adhesion and protein-protein interaction energies (green). The grey dashed line indicates the time when the cell-like vesicle detached from the target vesicle. This detachment allows the cell-like vesicle to retract its long thin protrusion, thereby reducing its bending energy.

**Functionalization of hydrogel beads with TDA**—Poly-Aam-co-AAc microparticles (DAAM particles, termed hydrogel beads here) were synthetized as described previously [14]. To functionalize them, 125$\mu$L of 5% v/v hydrogel beads were washed twice in activation buffer (100 mM MES (Sigma), pH 6.0, 200 mM NaCl) and subsequently incubated for 15 min in 125$\mu$L of activation buffer with 40 mg/mL 1-ethyl-3-(3-dimethylaminopropyl) carbodiimide (EDC, Sigma), 20 mg/mL N-hydroxysuccinimide (NHS, Sigma) and 0.1 % (v/v) Tween20 (Sigma). Next, the beads were centrifuged (1 min at 5000 g) and quickly resuspended in 125$\mu$L of 0.1 mg/mL Tetradecylamine (TDA, Sigma) dissolved in activation buffer. After 1 h incubation, the solution was centrifuged (1 min at 5000 g) to remove excess TDA and then resuspended with 125 $\mu$L of blocking buffer (300 mM Tris pH 9.0, 300 mM NaCl, 100 mM of Ethanolamine (Merck-Millipore). After 30 min incubation, functionalized beads were washed 3 times in 250 $\mu$L PBS with 0.1% (v/v) Tween20. Finally, hydrogel beads were resuspended in 125 $\mu$L PBS.

**Live cell in vivo microscopy**— Embryos were mounted in 1% low melting point agarose (UltraPure LMP Agarose, Invitrogen) in Danieau´s solution on a 35 mm glass bottom dish with 14 mm inner diameter of the glass surface (MatTek) and covered with Danieau´s solution. Embryos were imaged at Leica TCS SP8 or SP8 FALCON

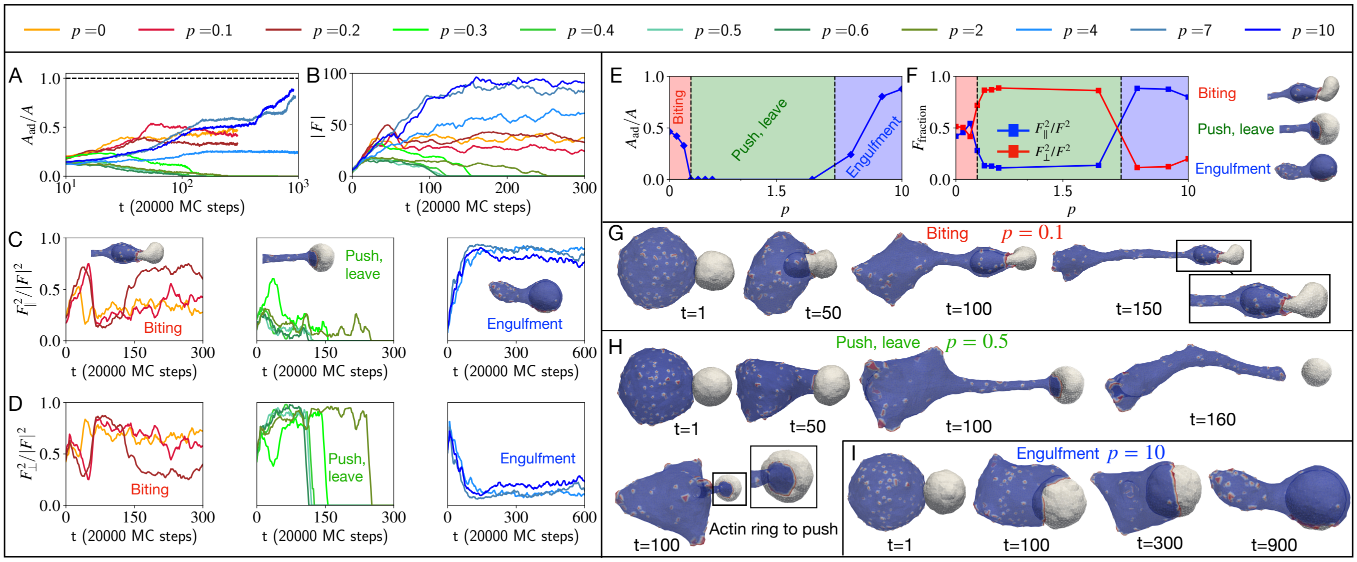


FIG. S9. Effect of the internal pressure of the target cell on the process of phagocytosis. The smaller vesicle (target) and the bigger vesicle (attacking cell) are made of 847 and 3127 vertices, respectively. The time evolution of the adhesive area fraction and magnitude of the force applied on the target cell by the attacking cell are shown in A) and B), respectively. C) It shows the fraction of tangential force on the target for three different cases of nibbling, push-leave, and engulfment in three different panels. D) Similarly, the fraction of normal or pushing force on the target for three different cases of nibbling, push-leave, and engulfment in three different panels. E) We have shown the steady state average value of adhesive area fraction with the bending rigidity $\kappa$ of the target cell. F) Here, we averaged the tangential and normal force fraction in the relevant time windows. (G-I) The time evolution of the shapes and the snapshots are shown for three different cases of nibbling, push-leave, and engulfment respectively.

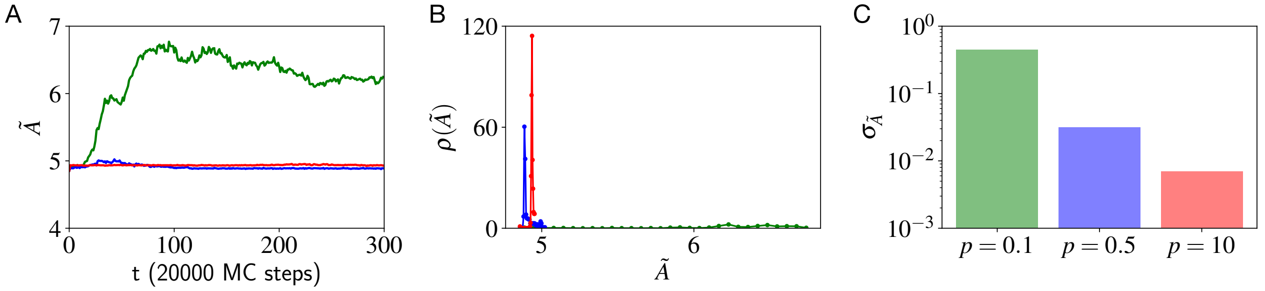


FIG. S10. A measure of membrane tension when the area and the volume both change over time during the active engulfment of a target-like vesicle with different internal pressure (Fig7G-I). A) The size normalised area $\tilde{A}$ (Eq.S12) for three different pressure differences $p = 0.1$, $p = 0.5$, and $p = 10$ shown in green, blue and red, respectively. B) The probability distribution of the size normalized area $\rho(\tilde{A})$. C) The standard deviation $\sigma_{\tilde{A}}$ of the size-normalized area. Higher tension results in lower standard deviation of the area fluctuations, and therefore higher internal pressure implies higher membrane tension. We set the bending rigidity of both the vesicles to $20k_BT$. The active force parameter $F = 2k_BT$, adhesion energy per node is $E_{\text{ad}} = 2k_BT$ between the two vesicles.

STED using a HC PL APO CS2 20 x/0.75 NA or HC PL APO CS2 40x/1.30 OIL immersion objectives. A laser excitation of 488 nm with a HyD or PMT detector and 561 nm with a HyD or PMT detector were used. For live imaging of apoptotic clearance and dispersal, z-stacks with a spacing of 2 $\mu$m between z-slices over a total depth of around 30 $\mu$m were acquired with a temporal resolution around 90 seconds. For high-resolution imaging of phagocytic cups 1 $\mu$m z-slices were acquired with a temporal resolution of 40 seconds. Embryos were imaged at 28°C using a H301-K stage top incubator (Okolab) with a UNO-T-H-CO2 controller (Okolab).

**Analysis of phagocytic efficiency of hydrogel beads** Quantification of phagocytic activity of hydrogel beads was performed using the 'Multi-point' tool in FIJI. Phagocytic efficiency was calculated as the ratio of the number of synthetic targets cleared by the embryonic epithelium over the total number of synthetic targets present in a given

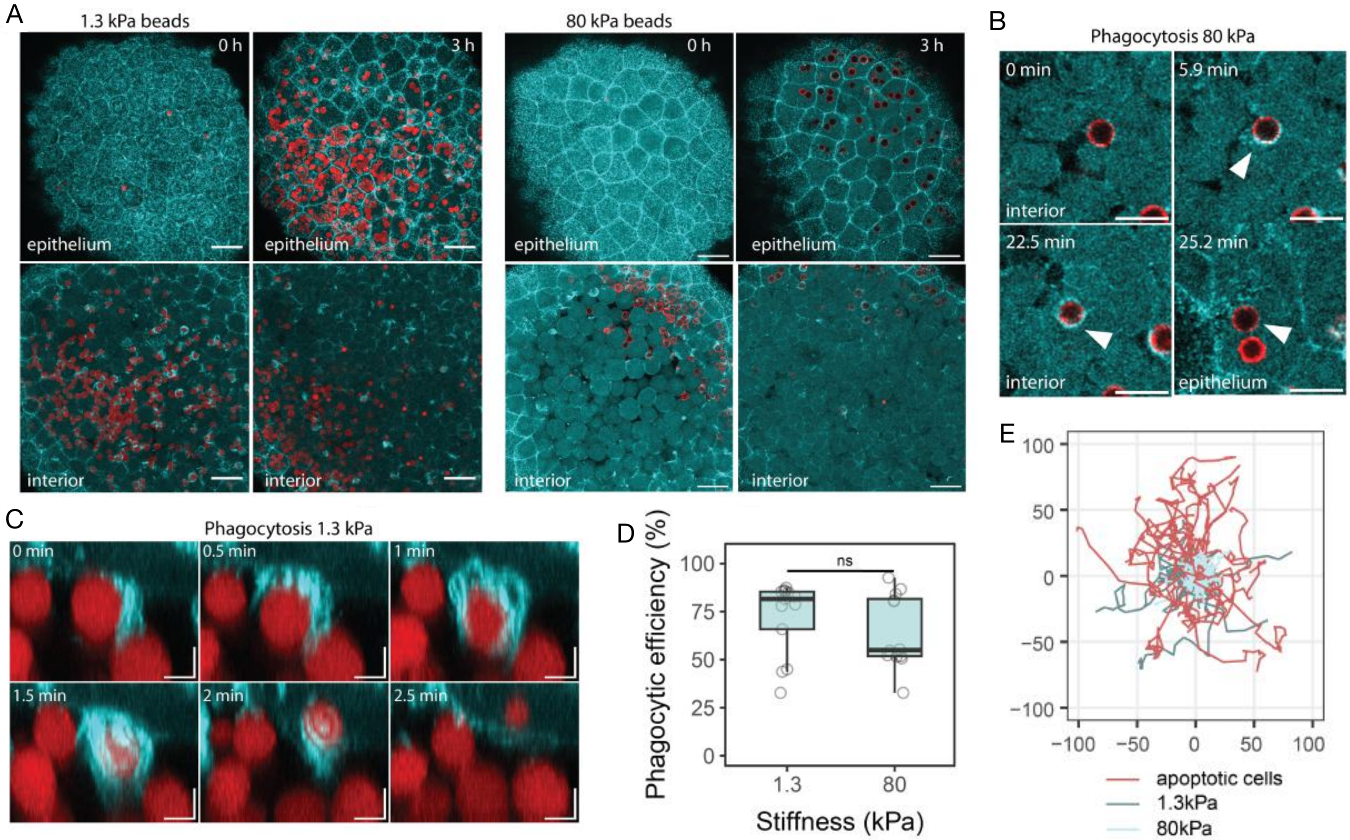


FIG. S11. **Epithelial phagocytosis and mobility of apoptotic cells and synthetic targets with variable target stiffness in vivo** — A) Representative images showing the presence of synthetic apoptotic targets (1.3 kPa, left; 80 kPa, right) in the interior of a zebrafish blastula stage embryo expressing Lifeact-GFP (cyan) at t=0 and their phagocytic clearance by the outer epithelial tissue over time (t=3 h). B) Representative images of an individual phagocytosis event of 80 kPa hydrogel beads in vivo. C) High-resolution transversal view of an individual phagocytic uptake event of 1.3 kPa hydrogel beads in a Lifeact-GFP (cyan) expressing embryo. D) Quantification of phagocytic efficiency derived as the percentage of synthetic targets cleared by the epithelium for 1.3 kPa hydrogel beads (n = 13 embryos) and 80 kPa hydrogel beads (n = 12 embryos). Data points represent the apoptotic target clearance efficiency as the percentage of cleared targets in individual embryos. N = 3 independent experiments. Welch Two Sample t-test, p = 0.99947. E. Tracks of individual apoptotic targets showing the path travelled over a maximum period of 30 min for apoptotic cells (red, n = 32 tracks from 3 embryos), 1.3 kPa hydrogel beads (dark cyan, n = 29 tracks from 3 embryos) and 80 kPa hydrogel beads (light cyan, n = 30 tracks from 3 embryos). Tracks were centered to the origin. Scalebars: 40$\mu$m (A), 20$\mu$m (B), 5$\mu$m z: 5$\mu$m (C).

field of view.

**Analysis of epithelial arm dynamics**— Tracking of apoptotic cells and synthetic targets was performed by using the 'MTrackJ' plugin [5]. A representative number of motile apoptotic or synthetic targets per embryo were tracked over time. The speed was calculated by the distance that targets moved in X-Y-Z directions between consecutive time frames (time lag tlag $\approx$ 90s). The maximum speed corresponds to the maximum instantaneous speed for each track. For the visualization of the spatial target spreading in the embryo in vivo, the x- and y- coordinates of targets were obtained over a time period of a maximum of 30 minutes. Tracks were aligned and centred at the origin.

**Movie S1: Passive engulfment of the soft vesicle:**— The cell-like vesicle with passive ($F = 0$) curved protein engulfing the target vesicle. We set the parameters for a cell-like vesicle $N = 3127$, $\kappa = 20\ k_BT$, $\rho = 4.8\%$, $c_0 = 1\ l_{\min}^{-1}$, $w = 1\ k_BT$. The parameters for a target vesicle are set as $N = 847$, $\kappa = 20\ k_BT$, $\rho = 0\%$. The adhesive energy between the cell-like vesicle and the target vesicle is $E_{\rm ad} = 2k_BT$.

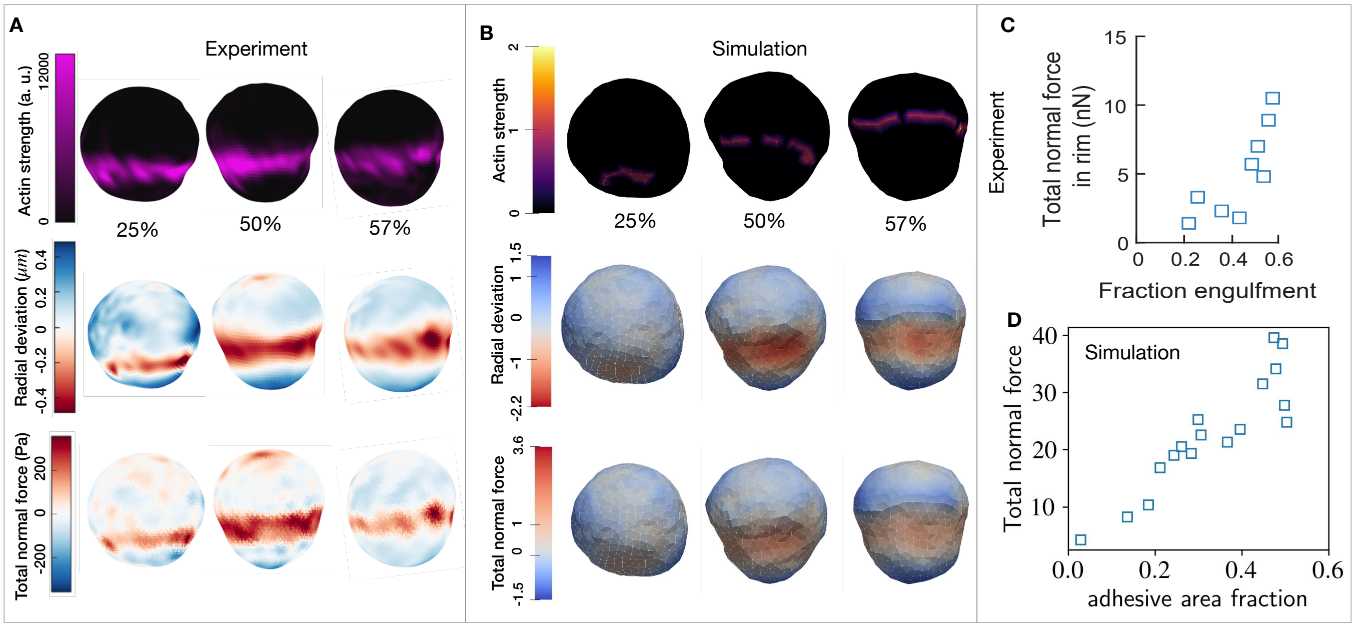


FIG. S12. A) 3D reconstructions of deformable acrylamide-co-acrylic acid-microparticles (DAAM-particles) (1.4 kPa, 9 μm) revealing F-actin over the particle surface, detailed target deformations induced during phagocytosis and normal forces inferred from shape deformations. DAAM-particles were functionalized with Immunoglobulin G (IgG) and stained with TAMRA-cadaverine and exposed to phagocytosis by RAW264.7 cells, which were then fixed and stained for F-actin. Each panel displays (left) F-actin distribution across the particle surface, (middle) radial deformation, and (right) the normal traction forces inferred from the shape deformations. (B) Snapshots of the simulated target vesicle at different values of engulfed fraction. Heatmaps show the location of the CMC (left), target vesicle radial deformation (middle) and the normal component of the active force exerted by the CMC on the target vesicle (right). These correspond to the actin localization in the experiments, bead deformation and normal elastic forces, respectively. In these simulations, we added the energy due to the bulk modulus of the target $\kappa_{bulk} = 0.5\ k_BT/l_{\min}^2$ (Eq.S13), while the bending modulus of the target is $\kappa = 20\ k_BT$. (C,D) plots of total normal force as function of the adhesive area fraction, in the experiment and simulation, respectively.

**Movie S2: (Experiment) GUV is engulfed by the macrophage**—Macrophages phagocytose taut GUVs. The macrophage cytosol is labeled by CellTracker Green CMFDA, and the actin is labeled by LifeAct GFP (green). The GUV is opsonized with fluorescent anti-biotin AlexaFluor 647 (magenta). Each frame is 30 seconds.

**Movie S3: (Experiment) GUV is pushed by the macrophage**—Macrophages push and then trogocytose low-tension GUVs. The macrophage cytosol is labeled by CellTracker Green CMFDA, and the actin is labeled by LifeAct GFP (green). The GUV is opsonized with fluorescent anti-biotin AlexaFluor 647 (magenta). Each frame is 30 seconds.

**Movie S4: (Experiment) GUV is bitten by the macrophage**—Macrophages trogocytose low-tension GUVs. The macrophage cytosol is labeled by CellTracker Green CMFDA, and the actin is labeled by LifeAct GFP (green). The GUV is opsonized with fluorescent anti-biotin AlexaFluor 647 (magenta). Each frame is 30 seconds.

**Movie S5: (Experiment) Lymphoma cell is engulfment by the macrophage**—The example of phagocytosis of a lymphoma cell (magenta) by a lifeact-expressing macrophage (green). Time stamp hours:minutes: seconds.

**Movie S6: (Experiment) Lymphoma cell is pushed by the macrophage**—The example of a lifeact-expressing macrophage (green) pushing a lymphoma cell (magenta). Time stamp hours:minutes: seconds.

**Movie S7: (Experiment) Lymphoma cell got bitten by the macrophage**—The example of trogocytosis/biting of a lymphoma cell (magenta) by a lifeact-expressing macrophage (green). Time stamp hours:minutes: seconds.

**Movie S8: Active cell-like vesicle engulfing the target**—The cell-like vesicle with active ($F \neq 0$) curved protein engulfing the target vesicle. We set the parameters for a cell-like vesicle $N = 3127,\ \kappa = 20\ k_BT,\ \rho = 4.8\%,\ c_0 = 1\ l_{\min}^{-1},\ w = 1\ k_BT,\ F = 2k_BTl_{\min}^{-1}$. The parameters for a target vesicle are set as $N = 847,\ \kappa = 20\ k_BT,\ \rho = 0\%$.

The target vesicle maintains an internal osmotic pressure $p = 10\ k_BTl_{\text{min}}^{-3}$. The adhesive energy between the cell-like vesicle and the target vesicle is $E_{\text{ad}} = 2k_BT$.

**Movie S9: Active cell-like vesicle pushing the target**—The cell-like vesicle with active ($F \neq 0$) curved protein pushing and finally leaves the target vesicle. We set the parameters for a cell-like vesicle $N = 3127$, $\kappa = 20\ k_BT$, $\rho = 4.8\%$, $c_0 = 1\ l_{\text{min}}^{-1}$, $w = 1\ k_BT$, $F = 2k_BTl_{\text{min}}^{-1}$. The parameters for a target vesicle are set as $N = 847$, $\kappa = 20\ k_BT$, $\rho = 0\%$. The target vesicle maintains an internal osmotic pressure $p = 0.5\ k_BTl_{\text{min}}^{-3}$. The adhesive energy between the cell-like vesicle and the target vesicle is $E_{\text{ad}} = 2k_BT$.

**Movie S10: Active cell-like vesicle biting the target**—The cell-like vesicle with active ($F \neq 0$) curved protein biting a portion of the target vesicle. We set the parameters for a cell-like vesicle $N = 3127$, $\kappa = 20\ k_BT$, $\rho = 4.8\%$, $c_0 = 1\ l_{\text{min}}^{-1}$, $w = 1\ k_BT$, $F = 2k_BTl_{\text{min}}^{-1}$. The parameters for a target vesicle are set as $N = 847$, $\kappa = 20\ k_BT$, $\rho = 0\%$. The target vesicle maintains an internal osmotic pressure $p = 0.1\ k_BTl_{\text{min}}^{-3}$. The adhesive energy between the cell-like vesicle and the target vesicle is $E_{\text{ad}} = 2k_BT$.

**Movie S11: Active cell-like vesicle biting the target instead of pushing, when held a patch**—The cell-like vesicle with active ($F \neq 0$) curved protein biting the target vesicle instead of pushing while held a circular patch by freezing (shown in black) the vertices of the target. We set the parameters for a cell-like vesicle $N = 3127$, $\kappa = 20\ k_BT$, $\rho = 4.8\%$, $c_0 = 1\ l_{\text{min}}^{-1}$, $w = 1\ k_BT$, $F = 2k_BTl_{\text{min}}^{-1}$. The parameters for a target vesicle are set as $N = 847$, $\kappa = 200\ k_BT$, $\rho = 0\%$. The adhesive energy between the cell-like vesicle and the target vesicle is $E_{\text{ad}} = 2k_BT$.

**Movie S12: In-vivo experiment**—In vivo tracking (yellow lines) of apoptotic targets (red, Bax+ cells co-expressing the plasma membrane marker Lyn-tdTomato, left), 1.3 kPa hydrogel beads (red, middle) and 80 kPa hydrogel beads (red, right) in Lifeact-GFP (cyan) expressing embryos. Embryos were obtained from the Tg(actb1:Lifeact-GFP) line. Time is indicated in h:min:s. Scale bar: 40 $\mu$m.

---


[1] C. E. Cornell, A. Chorlay, D. Krishnamurthy, N. R. Martin, L. Baldauf, and D. A. Fletcher. Target cell cortical tension regulates macrophage trogocytosis. *Nature Cell Biology*, pages 1–11, 2025.

[2] M. Fošnarič, S. Penič, A. Iglič, V. Kralj-Iglič, M. Drab, and N. S. Gov. Theoretical study of vesicle shapes driven by coupling curved proteins and active cytoskeletal forces. *Soft Matter*, 15(26):5319–5330, 2019.

[3] A. Mali, Y. Peeters, R. Rodrigues de Mercado, A. H. Settle, M. J. Footer, M. Srinivas, J. A. Theriot, and D. Vorselen. Using tunable hydrogel microparticles to measure cellular forces. *Nature Protocols*, Dec. 2025.

[11] D. Vorselen, S. Barger, J. Theriot, N. Gauthier, and M. Krendel. Phagocytic microscopy and mp-tfm assay with raw macrophages upon treatment with cytoskeletal inhibitors, 2021.

[12] D. Vorselen, S. R. Barger, Y. Wang, W. Cai, J. A. Theriot, N. C. Gauthier, and M. Krendel. Phagocytic 'teeth' and myosin-ii 'jaw' power target constriction during phagocytosis. *eLife*, 10, Oct. 2021.

[13] D. Vorselen, Y. Wang, M. M. de Jesus, P. K. Shah, M. J. Footer, M. Huse, W. Cai, and J. A. Theriot. Microparticle traction force microscopy reveals subcellular force exertion patterns in immune cell–target interactions. *Nature communications*, 11(1):20, 2020.

[14] D. Vorselen, Y. Wang, M. M. de Jesus, P. K. Shah, M. J. Footer, M. Huse, W. Cai, and J. A. Theriot. Microparticle traction force microscopy reveals subcellular force exertion patterns in immune cell–target interactions. *Nature Communications*, 11(1), Jan. 2020.
